# Stochastic modeling of the dynamics of *Salmonella* infection of epithelial cells

**DOI:** 10.1101/2023.04.02.535247

**Authors:** Jennifer Hannig, Alireza Beygi, Jörg Ackermann, Leonie Amstein, Christoph Welsch, Ivan Ðikić, Ina Koch

**Affiliations:** Cognitive Information Systems, Kompetenzzentrum für Informationstechnologie, Technische Hochschule Mittelhessen, Friedberg, Germany; Department of Molecular Bioinformatics, Institute of Computer Science, Goethe University Frankfurt, Frankfurt a. M., Germany; Department of Internal Medicine I, Goethe University Hospital, Frankfurt a. M., Germany; Institute of Biochemistry II, Goethe University Hospital, Frankfurt a. M., Germany

**Author notes:** Correspondence April 2, 2023. These authors contributed equally to this work (Jennifer Hannig, Alireza Beygi).

**Keywords:** *Salmonella* infection, host-pathogen dynamics, stochastic modeling, discrete-state continuous-time Markov process, Gillespie algorithm, Petri net, model systems

## Abstract

Bacteria of the *Salmonella* genus are intracellular pathogens, which cause gastroenteritis and typhoid fever in animals and humans, and are responsible for millions of infections and thousands of deaths across the world every year. Furthermore, *Salmonella* has played the role of a model organism for studying host-pathogen interactions. Taking these two aspects into account, enormous efforts in the literature are devoted to study this intracellular pathogen. Within epithelial cells, there are two distinct subpopulations of *Salmonella*: (i) a large fraction of *Salmonella*, which are enclosed by vacuoles, and (ii) a small fraction of hyper-replicating cytosolic *Salmonella*. Here, by considering the infection of epithelial cells by *Salmonella* as a discrete-state, continuous-time Markov process, we propose a stochastic model of infection, which includes the invasion of *Salmonella* into the epithelial cells by a cooperative strategy, the replication inside the *Salmonella*-containing vacuole, and the bacterial proliferation in the cytosol. The xenophagic degradation of cytosolic bacteria is considered, too. The stochastic approach provides important insights into stochastic variation and heterogeneity of the vacuolar and cytosolic *Salmonella* populations on a single-cell level over time. Specifically, we predict the percentage of infected human epithelial cells depending on the incubation time and the multiplicity of infection, an d the bacterial load of the infected cells at different post-infection times.

## 1 Introduction

The intake of contaminated food or water can be the cause of *Salmonella enterica* serovar Typhimurium—hereafter *Salmonella*—infection, which is responsible for yearly thousands of deaths around the world [1]. Infections with bacteria of the *Salmonella* genus lead to clinical syndromes ranging from mild gastroenteritis to severe enteric fever. Over recent decades, the increased occurrence of strains with resistance to antibiotics has become a serious health problem [2, 3], and urgent need demands for alternative and effective drugs against resistant intracellular bacteria. Bacteria of the *Salmonella* genus serve as a model organism to study the general mechanisms involved in bacterial infections [4, 5]. Extensive experimental studies have resulted in a deeper understanding of the underlying mechanisms of the invasion of epithelial cells by *Salmonella* [6–70]. *Salmonella* have been observed to swim close to the cell surface, the so-called near-surface swimming [48], where the bacteria can either take off from or dock to the cell surface via the needle-like structure of the type III secretion system (T3SS) encoded by *Salmonella* pathogenicity island 1 (SPI-1) [8, 14, 21, 23, 51]. The binding of *Salmonella* to the cell surface and the injection of the effector proteins trigger a signal in epithelial cells that causes the rearrangement of the cytoskeleton of the host cell, which results in a deformation of the cell membrane and the formation of the membrane ruffle around the docked bacterium [9, 17]. Such a ruffle on the cell surface forms an obstacle for other bacteria during their near-surface swimming. The frequent stop of bacteria at the ruffle leads to the trapping of more *Salmonella* close to the first docked bacterium. Thus, exploiting the preexisting ruffles, *Salmonella* invade epithelial cells in a cooperative manner [48, 57]. A vacuolar membrane, the *Salmonella*-containing vacuole (SCV) [11], surrounds the invading bacteria, where it also provides a replication niche, mediated by the SPI-2 [12, 13, 20, 23, 51]. The majority of the bacteria stay inside the protective enclosing of the SCV. However, in inbred mouse strains and tissue culture, it has been observed that a small fraction disrupts the membrane of the SCV and enters the host cell’s cytosol [20, 27, 30, 32, 39]. *Salmonella* with cytosolic access replicate at higher rates than those inside the SCV [27, 28, 39, 46, 55], with the risk that the host cells can counteract against the fast-replicating *Salmonella* by the antibacterial host defense, known as autophagy [35, 53, 56], also referred to as xenophagy [43]. With this mechanism, host cells can capture and eliminate intracellular pathogens as an innate immunity process [32, 62]. As well in mice as in cell culture, xenophagy has been reported to repress the dissemination of cytosolic *Salmonella* [32, 54]. At the later time point of infection, when the number of bacteria inside the epithelial cells becomes high, cells undergo inflammatory death and *Salmonella* are released into the extracellular microenvironment [46]. We have depicted the main stages of the invasion of epithelial cells by *Salmonella* in Fig. 1.

**Figure 1:**
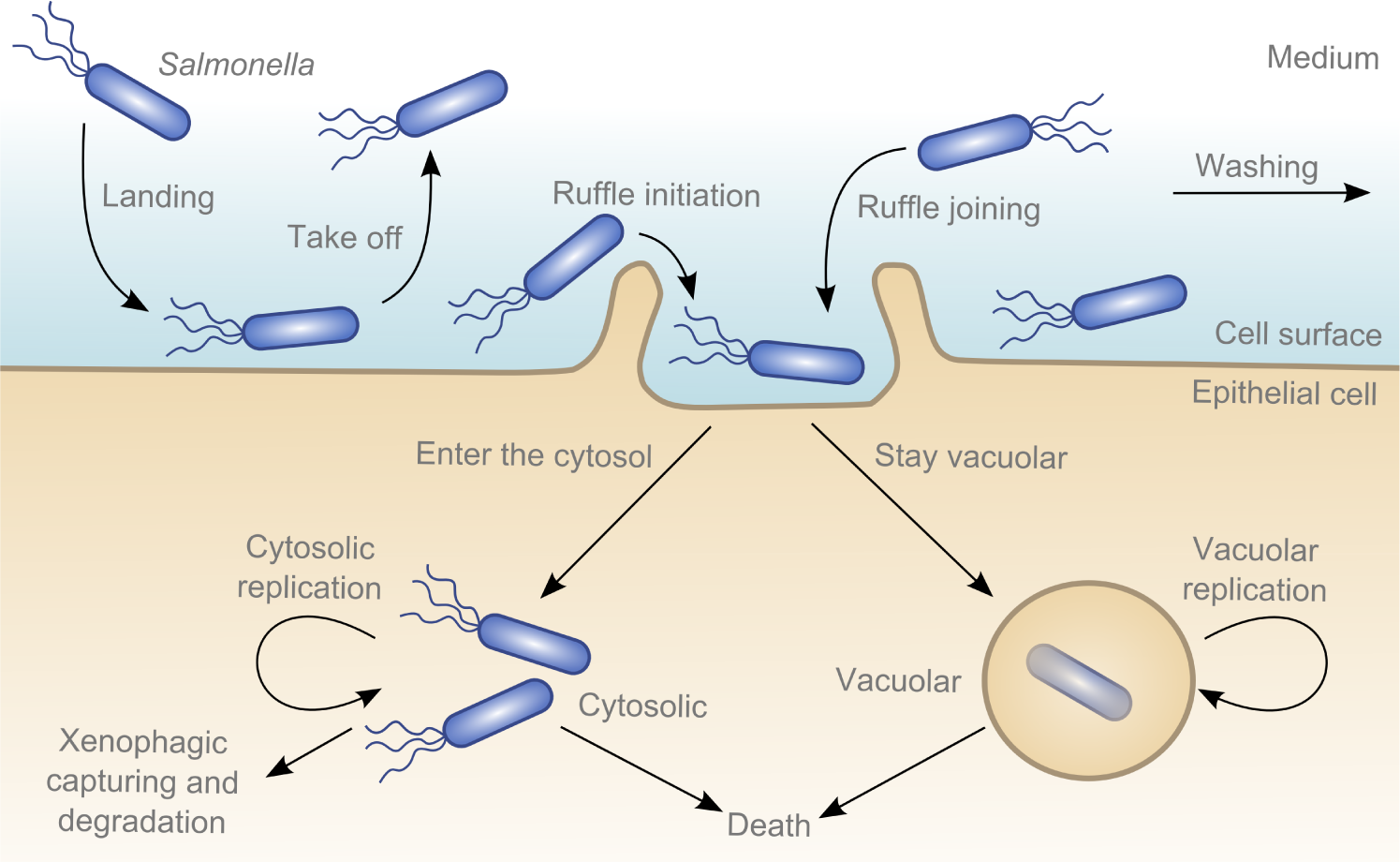
Schematic representation of the infection of the epithelial cell by *Salmonella*. Bacteria first land on the cell surface and perform near-surface swimming. Some take off and swim back to the medium and some dock to the cell surface. The binding of the bacterium results in *Salmonella*induced membrane ruffling, at which other bacteria could be trapped and joined during their near-surface swimming. In this way, bacteria cooperatively invade the host cell. Invasive *Salmonella* are enclosed within the SCV and start to replicate there. Some bacteria can escape from the SCV and enter the cytosol, where their replications are much faster. However, cytosolic *Salmonella* are vulnerable, as they can be recognized and degraded by xenophagy. Later in the infection process, cells that are populated by a large number of *Salmonella* undergo inflammatory death. In the experimental setting, the time period of infection ends with a washing step to remove the non-invaded *Salmonella*. The picture is adapted from [71].

On the theoretical side, several computational models have been proposed in the literature. For an overview of systems biological modeling of *Salmonella* metabolism, including the flux balance analysis and elementary flux modes approaches, see [45]. The communication between the three pathogenicity islands, T3SSs encoded by SPI-1, SPI-2, and T6SS, located in the genome of *Salmonella*, has been described as a Boolean network in [72]. A Petri net model of the xenophagic capturing of *Salmonella* in epithelial cells is presented in [73] to investigate the effects of *in silico* knockouts of autophagy receptors on the xenophagy pathway. For a summary of computational models concerning autophagy, we refer to [74]. In [75], a proteomics approach is presented, where it reveals dynamic changes in the global ubiquitinome of epithelial cells in response to infection by *Salmonella*. Kinetic models have been suggested to study different mechanisms of the accumulation of *Salmonella* in solid tumors [76] and to investigate the distribution of bacteria with respect to bacterial and cellular demography [77]. In the field of predictive microbiology, several quantitative models have been proposed to study microbial behavior, specifically the growth, in various foods [78–83]. Transmission dynamics of *Salmonella* Cerro in a US dairy herd is modeled in [84] via a set of nonlinear differential equations. Using continuous-time Markov chains [85, 86], dynamics of *Salmonella* infection are studied in UK dairy and pig herds. A stochastic model is constructed in [87] to investigate the dynamics of bacterial growth over the first few days of an acute *Salmonella* infection in mice. In [48], *Salmonella* are represented as linearly moving particles in a 3D landscape, consisting of a spherical object which is partially immersed into a flat surface. Simulation can show the accumulation of particles on the spherical object, whereas particles swim close to the surface (near-surface swimming). Invasion of macrophages by *Salmonella* is modeled in [88], where the probability of infection, after initial contact, is calculated to be remarkably low.

In this paper, we develop the first stochastic model for the infection of epithelial cells by *Salmonella*. The rationale behind considering the stochastic approach is the discreteness and the low number of bacteria that can attach to and invade a single epithelial cell. The low number of bacteria can cause significant stochastic effects and can lead to high heterogeneity in the bacterial population between individual cells, which becomes crucial for the prediction of the percentage of infected cells, i.e., the fraction of cells that harbor at least one bacterium, and the percentage of non-infected cells, i.e., the fraction of cells without bacteria inside. Deterministic models based on differential equations cannot capture the variability or random variation of the intracellular bacterial population. In the experimental studies, where the population number of invaded *Salmonella* are reported [41, 46, 48, 55], infected cells are monitored at a single-cell level, too.

We apply the Gillespie stochastic simulation algorithm to the known biological processes underlying the invasion of epithelial cells by *Salmonella*. The Gillespie algorithm is originally proposed in the context of chemical reactions [89, 90]. However, it has also been applied extensively in simulating stochastic processes in biology [91]. It presumes that the involved processes satisfy the Markov property, that is, the probability distribution of the future is only dependent on the present or the distribution is memoryless. Here, we assume the validity of the Markov property. We refer to [92] for a successful application of this algorithm in models of within-host pathogen growth in the case of a baculovirus of the gypsy moth.

This paper is organized as follows. In Sec. 2, we provide an overview of the underlying biological processes in the infection of epithelial cells by *Salmonella*. Exploiting the data reported in the literature, we estimate the parameters of our model and obtain the corresponding stochastic rate constants. In Sec. 3, we present two predictions of our model, that is, the percentage of infected cells, depending on the infection time and multiplicity of infection, and composition of the bacterial load. We conclude and summarize in Sec. 4.

## 2 Biological processes: stochastic modeling

In this section, we present the main features of our stochastic model, i.e., the biological processes that we have considered and the corresponding estimation of the model parameters based on the data reported in the literature. All essential features are summarized in Appendix. When the model is set up, we apply it to make testable predictions in the next section.

### 2.1 Development of the model

In Secs. 2.1.1 to 2.1.7, we describe the model development with the underlying biological processes. For a better understanding, we have depicted the underlying processes as Petri nets. A summary of the involved processes is given in Appendix A and Appendix B.

Petri net formalism is widely used for modeling and simulation of biological systems [93]. In a nutshell, Petri nets are weighted, directed, bipartite graphs, consisting of two types of vertices, places and transitions. Places depicted by circles are the passive part, such as chemical substances or biological organisms or species, and transitions depicted by rectangles are the active part, such as chemical reactions or biological processes. The dynamics of the Petri net is performed by the flow of discrete, movable objects, called tokens, denoted as *m*(·). Tokens are located on the places and move through the net via the activation of transitions. The initial marking of each place, denoted as *m*_0_(·), assigns a number of tokens to each place. Tokens can indicate, for example, the number of molecules or biological species. In the graphical representation, tokens are depicted by dots or numbers within places; an unmarked or empty place is indicative of zero tokens. Places and transitions are connected by arcs, which carry integer numbers corresponding to the arc weights; if no number is mentioned, the arc weight is one. These weights can represent the stoichiometric coefficients of the reactions or processes and thus determine the number of tokens that are consumed or produced by transitions. For a comprehensive overview of the Petri net approach in systems biology, we refer to [94].

#### 2.1.1 Generation of the infection medium

In the experimental setting, for the infection of cells by bacteria, often a particular multiplicity of infection (MOI) is indicated. The MOI is the ratio of bacteria in the infection medium to cells. For example, an indicated MOI of 100 in an experiment can be generated by 1000 bacteria in the infection medium to infect 10 cells. Thus, on average, 100 bacteria can potentially infect an individual cell. However, the actual MOI may differ due to uncertainties in the experimental setup. Also, the potential number of bacteria to infect an individual cell may vary from the indicated MOI due to an irregular distribution of the infection medium. To simulate these stochastic fluctuations, the generation of the infection medium is modeled by:

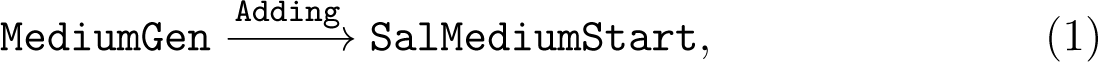

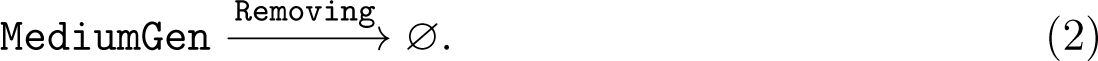

The initial marking of the place MediumGen indicates the maximum possible number of bacteria to infect an individual cell. The marking of the place SalMediumStart represents the actual number of *Salmonella* in the medium to infect an individual cell, and its initial value is zero. In this paper, we consider the maximum value of MOI as 2000, i.e., *m*_0_ (MediumGen) = 2000. Each token on the place MediumGen represents a bacterium that can either be added to the infection medium by the transition Adding or be removed by the transition Removing, where ∅ denotes the empty set. The transitions Adding and Removing are associated with the stochastic rate constants *c*_Adding_ and *c*_Removing_, respectively. In Sec. 2.2, we derive the corresponding rate constants. See Fig. 2, for the graphical representation of the processes (1) and (2).

**Figure 2:**
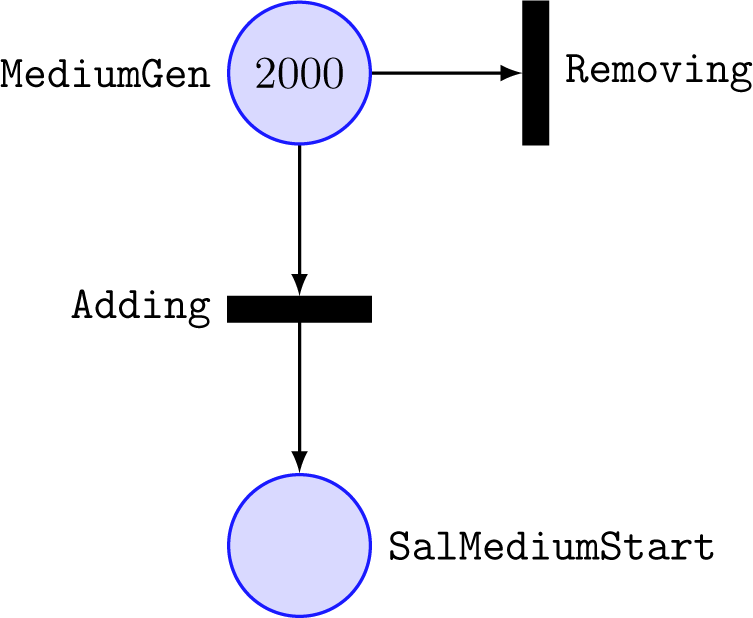
Symbolic representation of the generation of the infection medium. Circles (places) and rectangles (transitions) denote species and processes, respectively. The initial marking of the place MediumGen is shown as a number within the circle. The empty place SalMediumStart indicates zero tokens on that. Since the stoichiometric coefficients in the processes (1) and (2) are equal to one, the arc weights are set to one, indicated by no label.

The probability distribution of the number of bacteria to infect an individual cell is given by the binomial distribution,

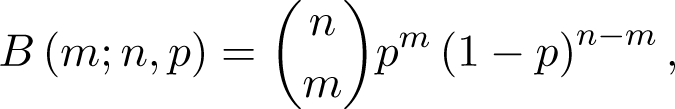

where *m* = number of bacteria per individual cell, *n* = *m*_0_ (MediumGen), and *p* = *c*_Adding_*/*(*c*_Adding_ + *c*_Removing_). The MOI is given by the mean of the binomial distribution, that is,

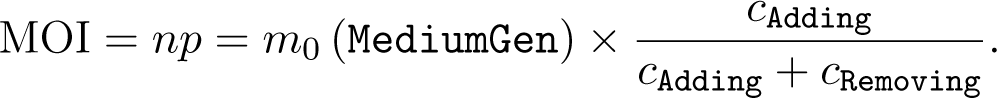

#### 2.1.2 *Salmonella* at different stages of the infection Landing and take off

We assume that *Salmonella* need some time to adjust to the medium at the beginning of the infection process. To model this adjustment, we have distinguished between the number of *Salmonella* in the medium at the beginning of the infection process, which is represented by the place SalMediumStart, and the number of the adapted bacteria to the medium, which is represented by the place SalMedium. The adaptation is achieved after the first landing of the bacteria on the cell surface and the subsequent taking off. First, we model the process of the initial landing of the bacterium on the cell surface as

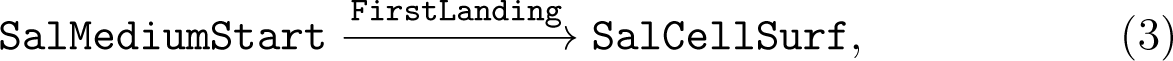

where the place SalCellSurf denotes the number of bacteria on the cell surface, with *m*_0_ (SalCellSurf) = 0. After some time of near-surface swimming, the bacterium takes off from the cell surface:

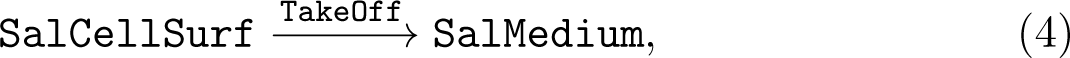

with *m*_0_ (SalMedium) = 0. By taking off from the cell surface, the bacterium goes back to the medium. This bacterium is now considered to be adapted to the infection medium. Landing of the adapted *Salmonella* on the cell surface is modeled by:

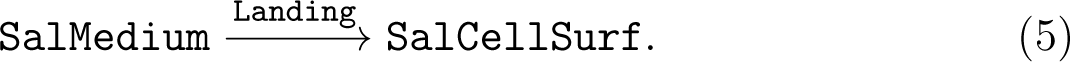

##### Ruffle initiation and joining

The formation of a ruffle on the cell surface is modeled by:

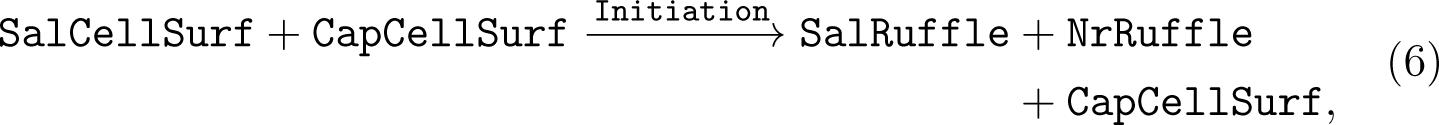

with *m*_0_ (CapCellSurf) = 1, *m*_0_ (SalRuffle) = 0, and *m*_0_ (NrRuffle) = 0. The place CapCellSurf describes a signal for the capability of the cell surface to accommodate additional bacteria, the place SalRuffle denotes the total number of bacteria located in ruffles, and the place NrRuffle counts the number of ruffles on the cell surface. For *m*_0_ (CapCellSurf) = 1, there is sufficient space on the cell surface to form a new ruffle by the transition Initiation. At each time when a new ruffle is formed, NrRuffle increases by one. We consider the attachment of *Salmonella* to the cell surface via the SPI-1-encoded T3SS to be irreversible [40]. Thus, in our model, we have accomplished that once a bacterium is located in a ruffle, it never leaves the ruffle again.

We model the cooperative invasion of *Salmonella* by exploiting the preexisting ruffles as

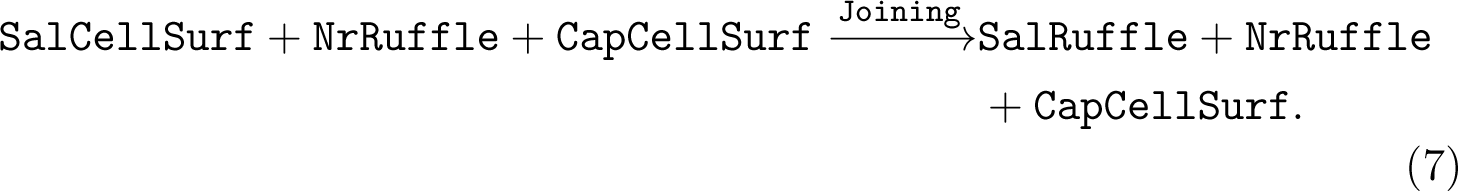

If at least one ruffle has been formed on the cell surface, i.e., *m* (NrRuffle) ≥ 1, a bacterium can join the ruffle by the transition Joining. For *m*(CapCellSurf) = 1, the cell surface has sufficient space for *Salmonella* to join the preexisting ruffles.

Due to the physical limitation of the cell surface, overall only a few ruffles can be formed. We model this restriction on the maximum number of ruffles on the cell surface as

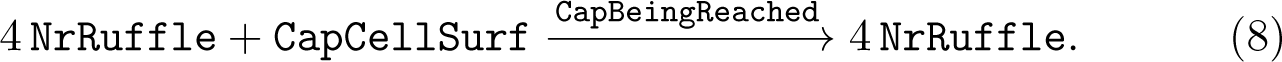

If the number of ruffles reaches four, i.e., *m* (NrRuffle) = 4, the transition CapBeingReached takes place, and the marking of the place CapCellSurf drops to zero. For *m* (CapCellSurf) = 0, no additional bacteria can be accommodated on the cell surface.

##### Staying vacuolar and entering the cytosol

Inside the cell, the decision of the bacterium, to stay either in the vacuole or enter the cytosol, is not entirely independent from the host cell, but it is also a cell-dependent process [95]. Here, we assume that there are cells with the capability to host *Salmonella* in their cytosol called cytosolic-capable cells and others are cytosolic-incapable. It has also been observed that cells containing cytosolic *Salmonella* can additionally contain vacuolar bacteria [55]. Thus, in cells with the ability to harbor cytosolic *Salmonella*, the bacterium can still decide whether to stay in the vacuole or go into the cytosol.

The process of staying of *Salmonella* in the SCV is modeled by

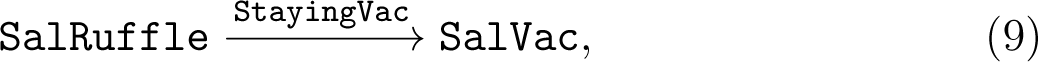

with *m*_0_ (SalVac) = 0. The place SalVac denotes the total number of bacteria in the SCV. The process of entering of *Salmonella* into the cytosol is modeled by

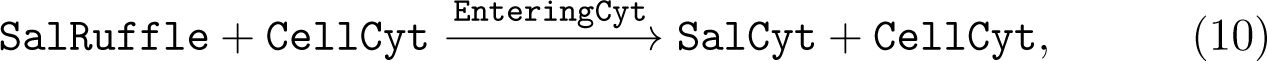

with *m*_0_ (CellCyt) = 0 and *m*_0_ (SalCyt) = 0. The place CellCyt describes a signal for the capability of the cell to host cytosolic bacteria and the place SalCyt represents the total number of *Salmonella* in the cytosol. Only if *m* (CellCyt) = 1, the transition EnteringCyt can occur and produces tokens on the place SalCyt. In other words, only if the cell is cytosolic-capable, the bacterium can enter the cytosol via the transition EnteringCyt. For simplicity, we have not included an intermediate step, where the bacterium is first located in a damaged SCV and, then, enters cytosol. In our model, *Salmonella* are directly translocated from the ruffle to the cytosol. This is justified as the vacuolar escape seems to be an early event, which happens soon after the internalization [55], and the bacteria within the SCV with the damaged membrane can be considered as a transient state between the initial intact SCV and the bacteria in the cytosol [60].

Overall, there are various scenarios for the cytosolic-capable cells: cells with exclusively *Salmonella* in the SCV, cells with exclusively *Salmonella* in the cytosol, cells with cytosolic and vacuolar *Salmonella*, and uninfected cells without invading bacteria. Cells with exclusively *Salmonella* in the SCV or uninfected cells can also originate from the cytosolic-incapable cells, where *m* (CellCyt) = 0.

##### Bacterial proliferation

The proliferation of *Salmonella* in the SCV and cytosol are modeled by the second-order autocatalytic processes:

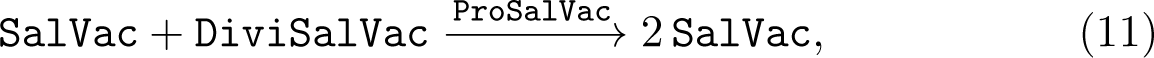

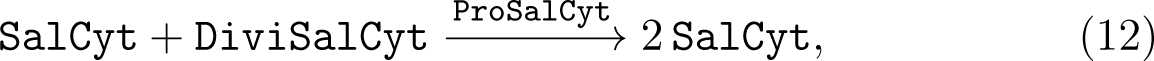

where the places DiviSalVac and DiviSalCyt, with *m*_0_(DiviSalVac) = *m*_0_(DiviSalCyt) = 0, model the lag and the stationary phases of the vacuolar and cytosolic replications, respectively. Each time the transition ProSalVac takes place, a token is removed from the places and two tokens are produced on the place SalVac, that is, one bacterium in the vacuole multiplies to two bacteria. In the same way, the transition ProSalCyt represents the proliferation of cytosolic *Salmonella*. In Fig. 3, we have illustrated the crucial steps in the infection of epithelial cells by *Salmonella*, described in the processes (3) to (12).

**Figure 3:**
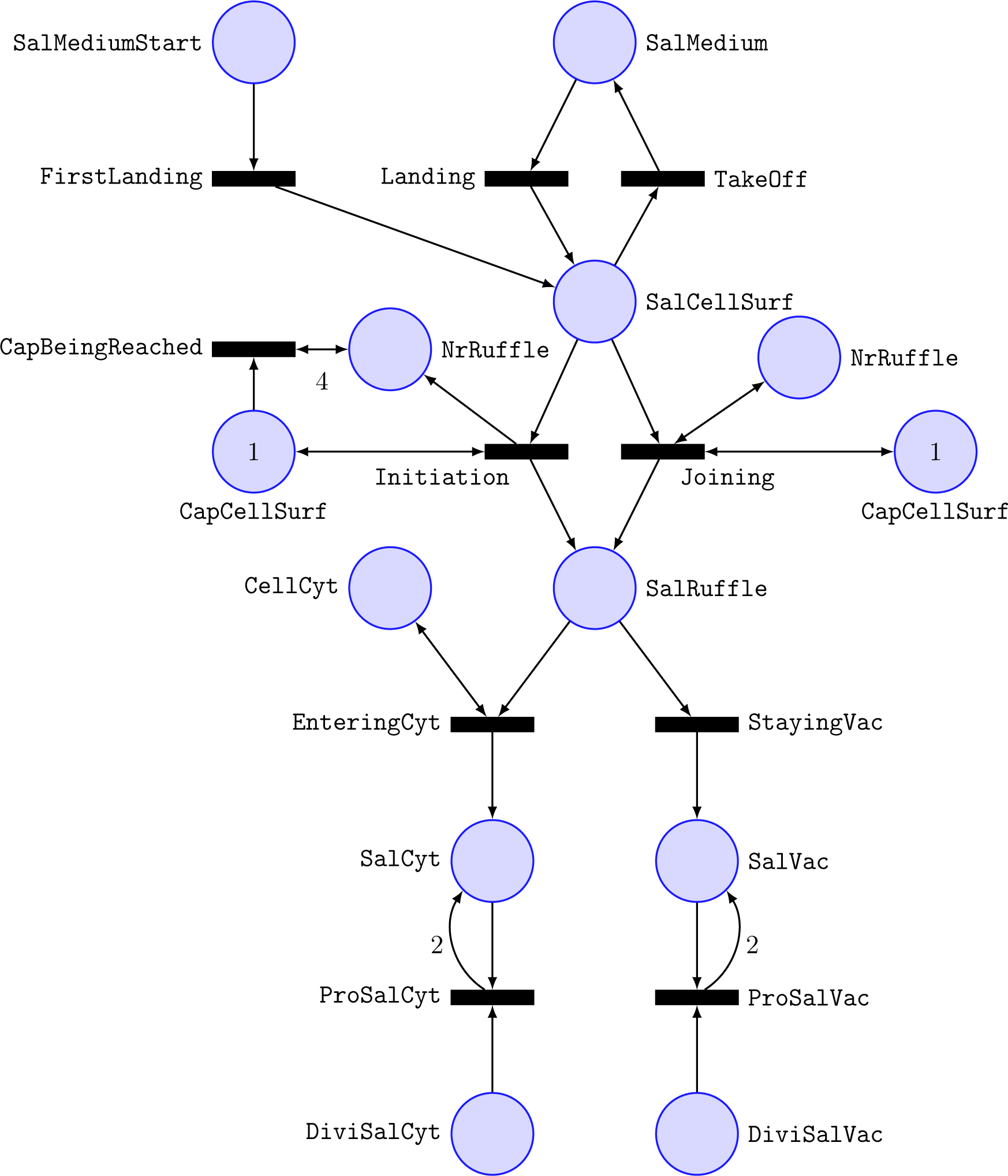
Different stages of the infection of the epithelial cell by *Salmonella*. Circles and rectangles denote species and processes in (3) to (12). Numbers within circles show the initial markings of species (no number implies zero) and the arc weights represent the stoichiometric coefficients (no number implies one).

#### 2.1.3 Fate of the epithelial cell

To model the cellular decision-making, whether or not a cell makes a choice to host *Salmonella* in its cytosol, we consider:

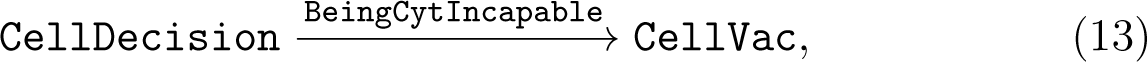

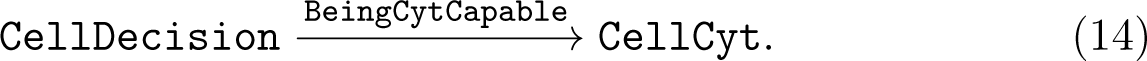

The place CellDecision, with *m*_0_ (CellDecision) = 1, represents the cell before making a choice. Occurrence of the transition BeingCytIncapable or BeingCytCapable produces a token on the place CellVac, where *m*_0_(Cell-Vac) = 0, or on CellCyt. When *m* (CellVac) = 1, cell is cytosolic-incapable and for *m* (CellCyt) = 1, cell is cytosolic-capable. See Fig. 4, for the symbolic representation of the processes (13) and (14).

**Figure 4:**
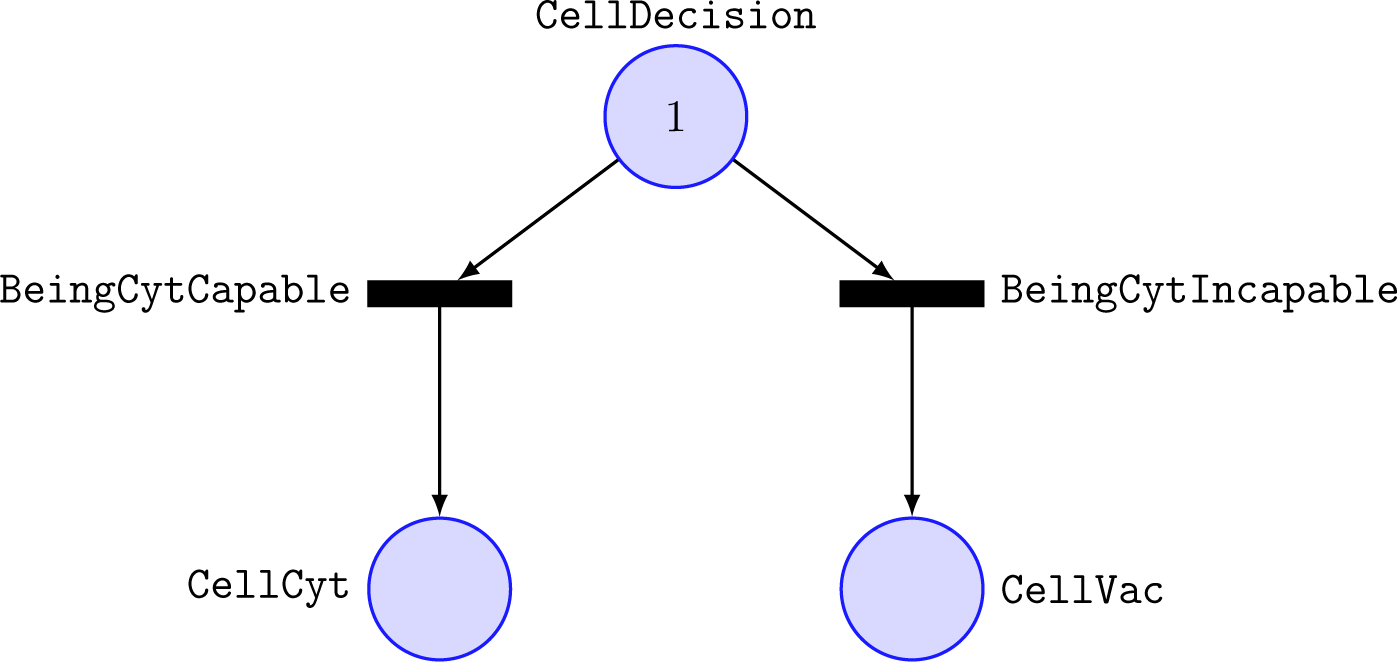
Graphical representation of cellular decision-making, where the cell makes a choice whether to become cytosolic-capable or cytosolic-incapable. Circles and rectangles correspond to the species and processes in (13) and (14). The initial marking of the place CellDecision is shown as a number within the circle.

#### 2.1.4 Divisibility of *Salmonella*

The doubling time of bacteria is dependent on their environmental conditions, such as space, nutrients, pH values, and temperature [96]. A change in the environmental conditions is followed by a time period during which bacteria need to adapt to their new environment called lag phase. During the lag phase, bacterial proliferation is impaired. Later in the infection process, the proliferation decreases due to limited nutrients and space called stationary phase. We combine the effects of the lag and stationary phases on bacterial growth into a feature, which we call divisibility. The divisibility of cytosolic and vacuolar *Salmonella* are represented by the places DiviSalCyt and DiviSalVac, respectively. In the following, we explain the underlying processes concerning the divisibility of cytosolic *Salmonella*. The case of vacuolar *Salmonella* is modeled in the same way.

The adaptability of *Salmonella* to the cytosol is denoted by the place AdapSalCyt, which can be increased by the processes:

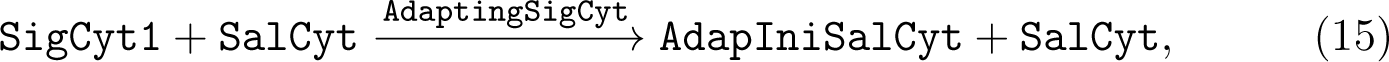

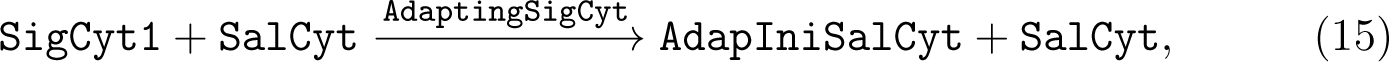

with *m*_0_ (SigCyt1) = 1, *m*_0_ (AdapIniSalCyt) = 0, and *m*_0_ (AdapSalCyt) = 0. The place SigCyt1 indicates a signal for the presence of *Salmonella* in the cytosol, the place AdapIniSalCyt represents a signal for the initiation of the adaptation process of *Salmonella* to the cytosol, and the place AdapSalCyt denotes the adjustment to the cytosol. If the bacterium enters the cytosol, i.e., *m* (SalCyt) ≥ 1, a token on the place SigCyt1 is consumed and produced on the place AdapIniSalCyt by the transition AdaptingSigCyt. For *m* (AdapIniSalCyt) = 1, the adaptability, AdapSalCyt, increases by the transition AdaptingCyt. Each time a token is produced on the place AdapSalCyt, it is consumed and produced on the place DiviSalCyt as

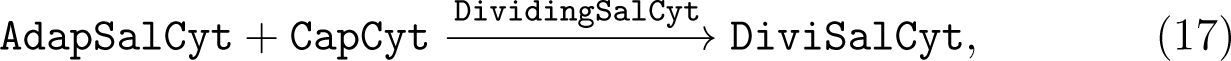

where the place CapCyt represents the capacity of the cytosol. To compute this capacity in the case of HeLa cells, first, we consider the HeLa cell volume as 2600 µm^3^ [98]. *Salmonella* have been reported to be 2 to 5 µm in length, with a diameter in the range of 0.7 to 1.5 µm [52]. Since *Salmonella* is rod-shaped, its average volume can be calculated as *V* = *πr*^2^*h* ≈ 3.33 µm^3^, where *r* = 0.55 µm and *h* = 3.5 µm. Thus, theoretically, up to 782 bacteria can fit into one HeLa cell. Due to cellular structure and geometrical considerations, the theoretical value is overestimated; instead, we assume the accommodation of up to 700 bacteria in one HeLa cell, i.e., *m*_0_ (CapCyt) = 700. Additionally, the place CapCyt can be interpreted as a measure for restrictions of environmental sources, e.g., space or nutrients. As the maximum divisibility of cytosolic *Salmonella* is dependent on the initial marking of the place CapCyt, we have Max (DiviSalCyt) = 700.

For *m* (DiviSalCyt) = 700, *Salmonella* proliferate with the highest achievable doubling time,

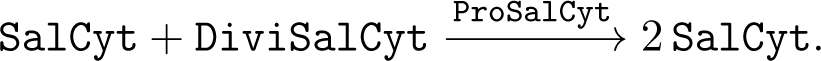

Each time the bacterium multiplies, a token on the place DiviSalCyt is consumed, which implies a decrease in the doubling time due to lack of nutrients or space. This situation resembles the stationary phase of the growth, as the number of cytosolic *Salmonella* converges towards 700.

Similar to the cytosolic case, we can model the divisibility of *Salmonella* in the SCV as follows:

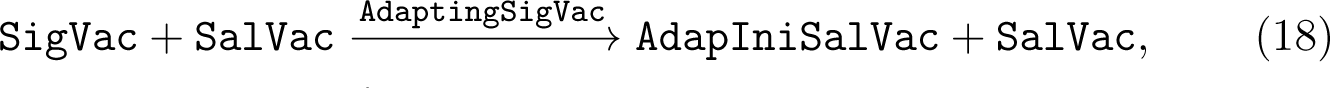

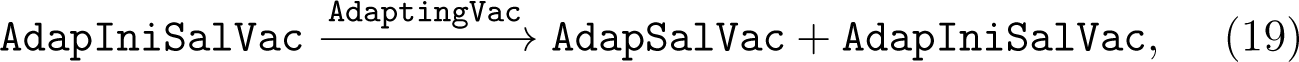

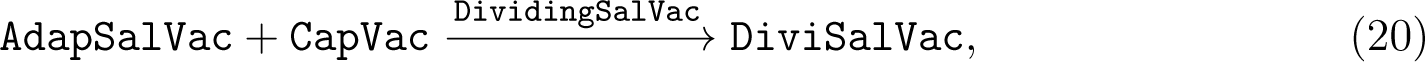

where *m*_0_ (SigVac) = 1, *m*_0_ (AdapIniSalVac) = 0, and *m*_0_ (AdapSalVac) = 0. We assume *m*_0_ (CapVac) = 150, that is, up to 150 bacteria can be accommodated in the SCV. See Fig. 5, for the depiction of the processes (15) to (20).

**Figure 5:**
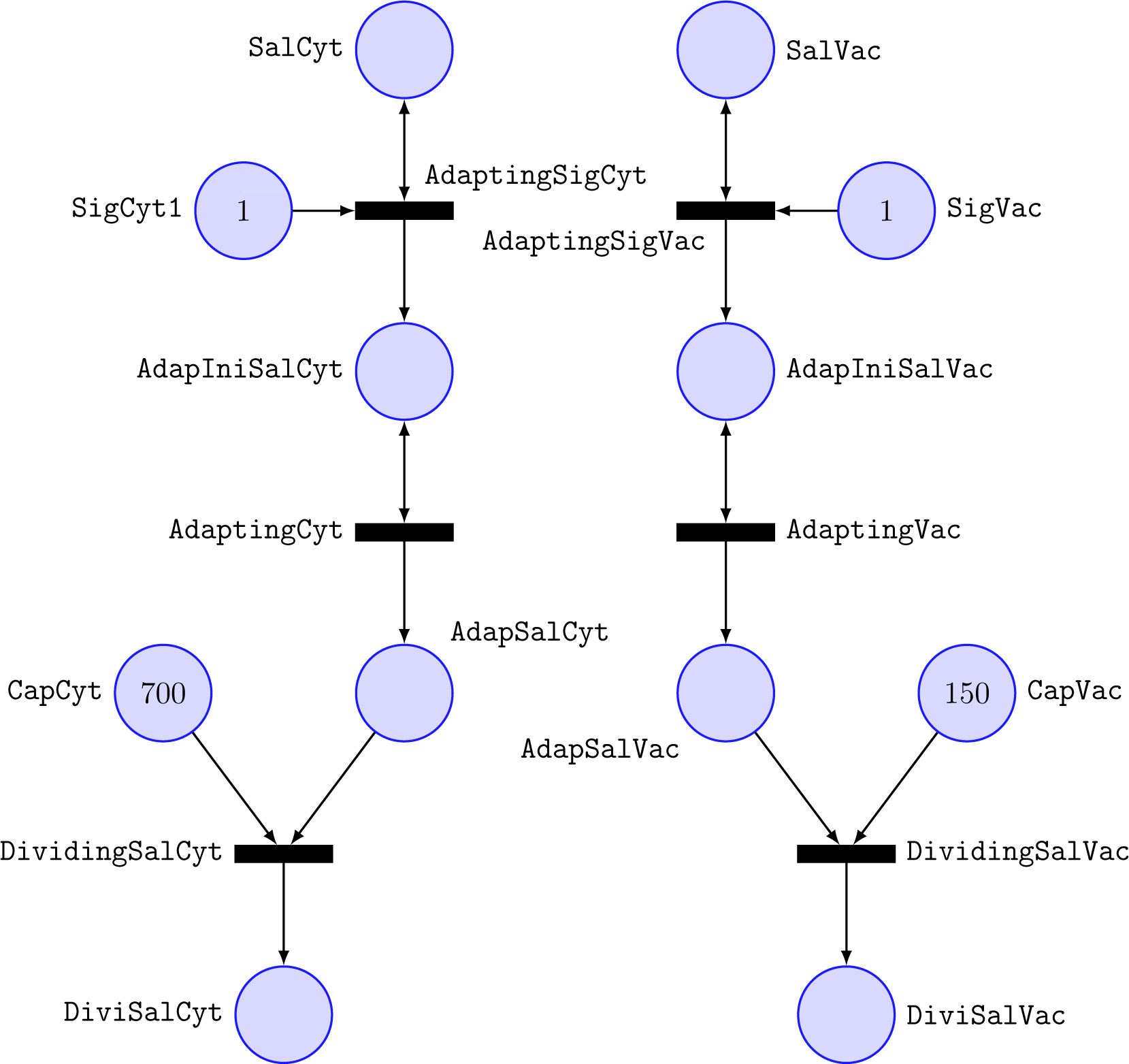
Graphical representation of (15) to (20), which model the effects of the lag and stationary phases on bacterial growth. Circles and numbers denote species and the corresponding initial markings, respectively, and rectangles represent processes.

#### 2.1.5 Xenophagic capturing and degradation of *Salmonella*

Degradation of cytosolic *Salmonella* by xenophagy is modeled by:

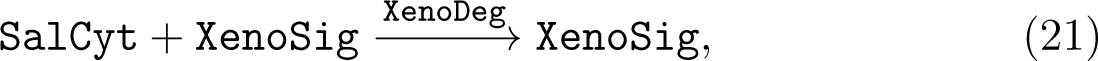

with *m*_0_ (XenoSig) = 0. The place XenoSig represents a signal for xenophagic degradation, which is activated when *m* (XenoSig) = 1. For *m* (XenoSig) = 1, tokens on the place SalCyt are consumed through the transition XenoDeg, i.e., bacteria are eliminated from the cytosol by xenophagic degradation.

It is suggested that xenophagy captures cytosolic *Salmonella* only in the early stages of the infection process [32, 47, 50], and after some time the autophagic targeting is prevented by the reactivation of mTOR [47, 50]. To take into account the temporary nature of the xenophagic recognition of cytosolic *Salmonella*, first, we consider the following process, which induces the initiation and termination of xenophagy:

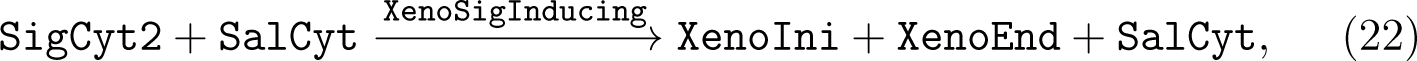

with *m*_0_ (SigCyt2) = 1, *m*_0_ (XenoIni) = 0, and *m*_0_ (XenoEnd) = 0. The places SigCyt2, XenoIni, and XenoEnd represent signals for the presence of *Salmonella* in the cytosol, the initiation, and the termination of xenophagy. If the bacterium reaches the cytosol, i.e., *m* (SalCyt) ≥ 1, a token on the place SigCyt2 is consumed by the transition XenoSigInducing and is produced on the places XenoIni and XenoEnd, i.e., *m* (XenoIni) = *m* (XenoEnd) =

## 1. In a nutshell, signals for the initiation and termination of xenophagy are induced

Xenophagy is initiated by:

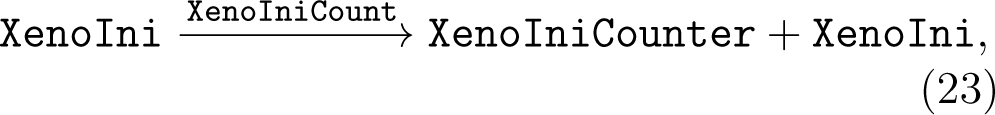

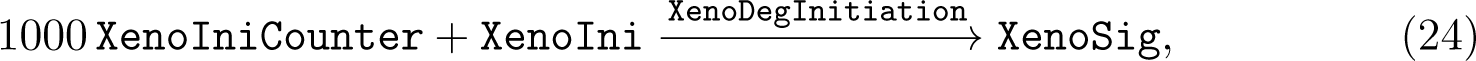

with *m*_0_ (XenoIniCounter) = 0, where the place XenoIniCounter denotes a time counter. For *m* (XenoIni) = 1, tokens are accumulated on the place XenoIniCounter by the transition XenoIniCount. If the time counter, the place XenoIniCounter, reaches 1000 tokens, the transition XenoDegInitiation generates one token on the place XenoSig, implying the activation of xenophagic degradation. Additionally, through this transition, the marking of the place XenoIni drops to zero, which in turn, stops the time counter and prevents further xenophagy initiation.

We can model the xenophagy termination in the same way:

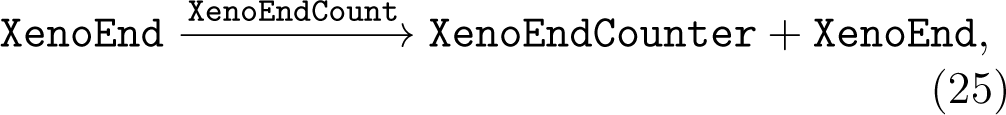

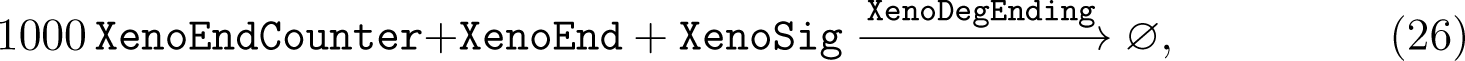

with *m*_0_ (XenoEndCounter) = 0. We have illustrated different stages of the xenophagic capturing and degradation of cytosolic *Salmonella*, i.e., the processes (21) to (26), in Fig. 6.

**Figure 6:**
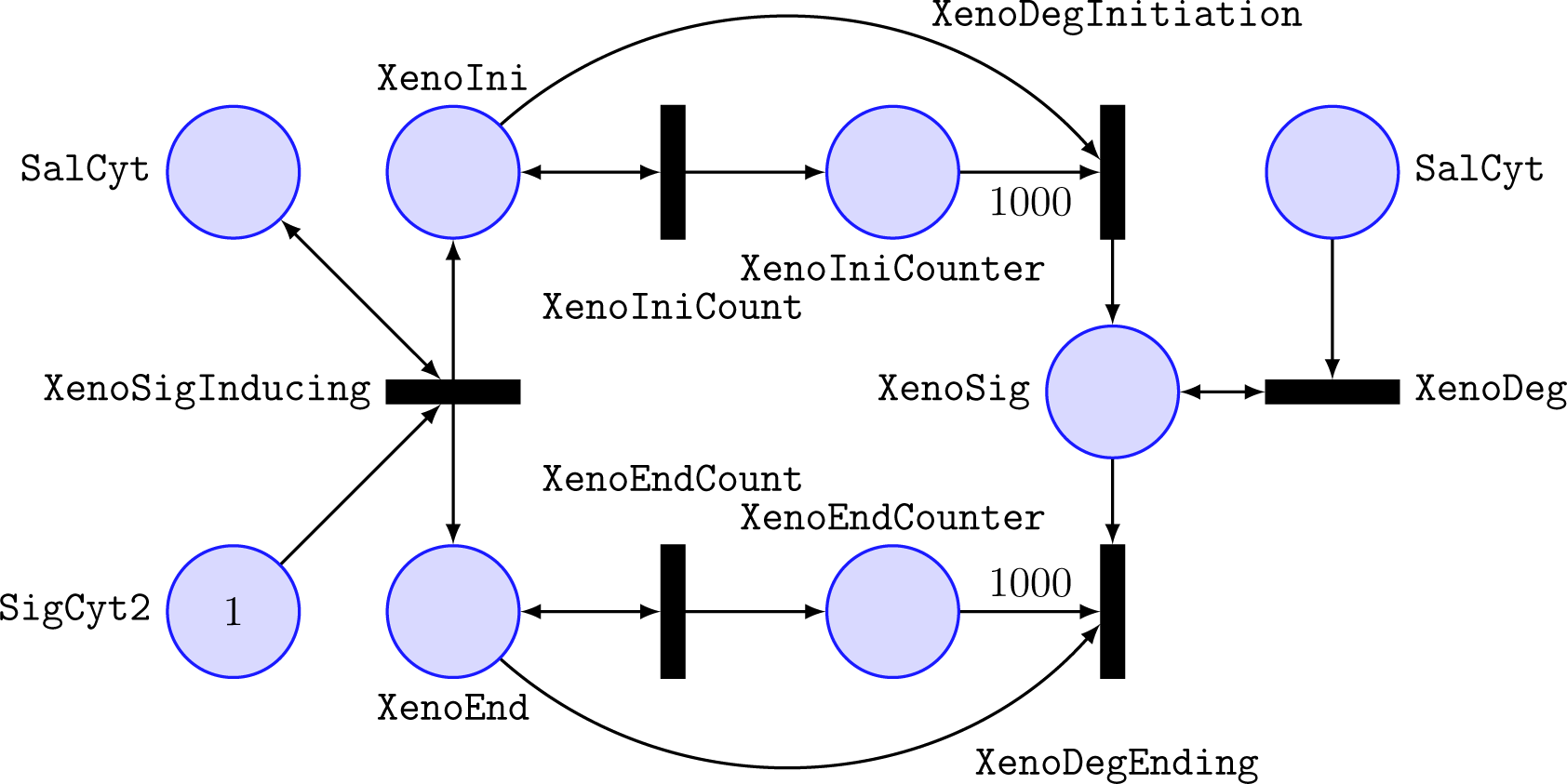
Recognition and elimination of cytosolic *Salmonella* by xenophagy. Circles and rectangles represent species and processes in (21) to (26). The initial marking of the place SigCyt2 is shown as a number within the circle and the arc weights 1000 correspond to the stoichiometric coefficients in (24) and (26).

### 2.1.6 Cell death

To model the inflammatory death, i.e., pyroptosis, of infected cells by cytosolic *Salmonella* [46, 58], we consider:

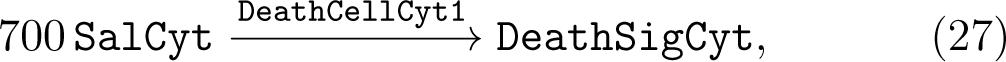

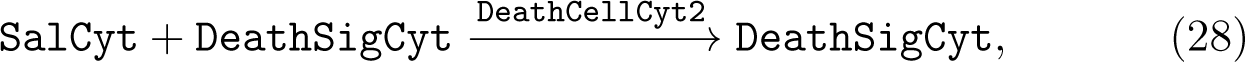

with *m*_0_ (DeathSigCyt) = 0. The place DeathSigCyt represents a signal that the cell undergoes inflammatory death due to a high number of cytosolic *Salmonella*. If the number of *Salmonella* in the cytosol reaches 700, i.e., *m* (SalCyt) ≥ 700, which is the upper limit of the number of bacteria that can be accommodated in the cytosol in our model, 700 tokens on the place SalCyt are consumed by the transition DeathCellCyt1. The remaining bacteria in the cytosol are removed by the transition DeathCellCyt2.

In the same way, we can model the death of cells by vacuolar *Salmonella*, due to the collapse of vacuolar membrane and triggering pyroptosis, as

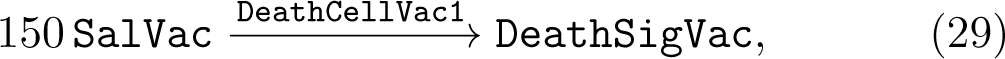

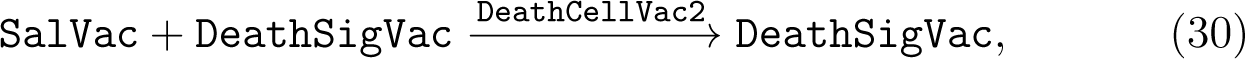

where the place DeathSigVac, with *m*_0_ (DeathSigVac) = 0, denotes a signal for cell death in this case.

For cells containing both cytosolic and vacuolar *Salmonella*, we consider:

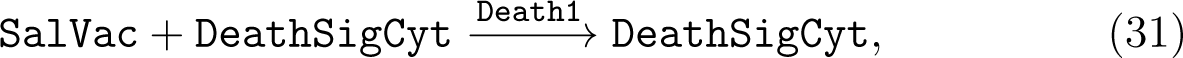

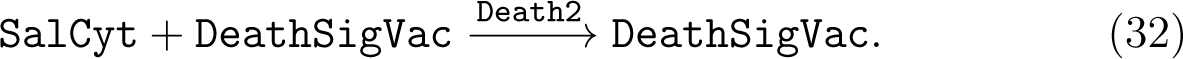

The above two processes remove vacuolar bacteria in cells that die as a result of a high number of cytosolic bacteria and remove cytosolic bacteria in cells that die as a result of a high number of vacuolar bacteria, respectively. In our model, we consider cell death as a process that happens at the later time point of infection, when the number of intracellular bacteria is high, as has been observed in [46]. See Fig. 7, for the illustration of the processes (27) to (32).

**Figure 7:**
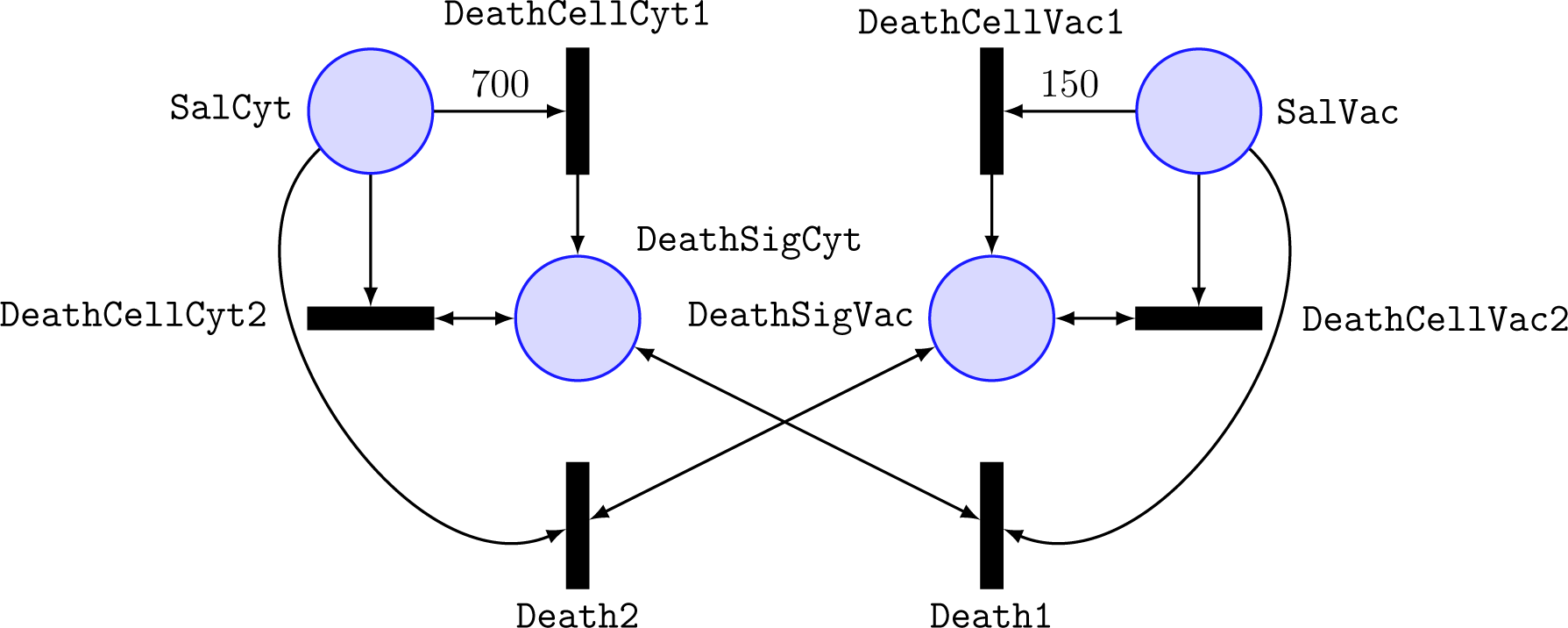
Symbolic representation of (27) to (32), which model *Salmonella*- induced, inflammatory cell death. Circles and rectangles denote species and processes, respectively. The weights of the arcs correspond to the stoichiometric coefficients.

### 2.1.7 Washing of the epithelial cell

We conclude Sec. 2.1 by incorporating a technical step which is often done in the experiments. The infection-time period in the experimental setting often ends with a washing step to remove the non-invaded *Salmonella*. We model the initiation of the washing process as

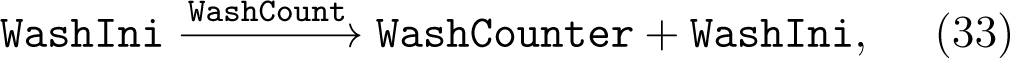

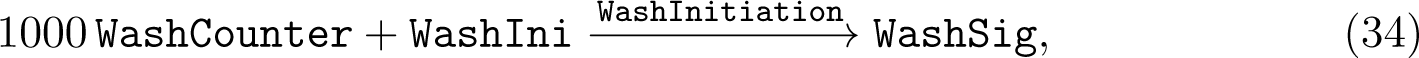

with *m*_0_ (WashIni) = 1, *m*_0_ (WashCounter) = 0, and *m*_0_ (WashSig) = 0. The places WashIni, WashCounter, and WashSig represent a signal for the one-time initiation of the washing process, a time counter, and a signal for the onset of the washing process, respectively. For *m*_0_ (WashIni) = 1, tokens are accumulated on the place WashCounter by the occurrence of the transition WashCount. When the place WashCounter reaches 1000 tokens, the transition WashInitiation generates one token on the place WashSig, which drops the marking of the place WashIni to zero, i.e., *m* (WashIni) = 0, preventing the initiation of the further washing process.

For *m*_0_ (WashSig) = 1, the washing process starts and the non-invaded bacteria in the infection medium or on the cell surface are removed by:

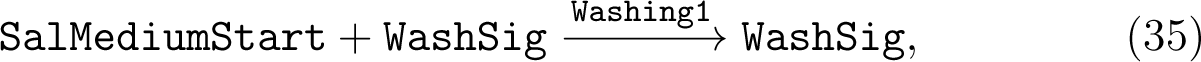

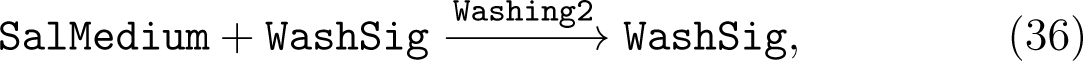

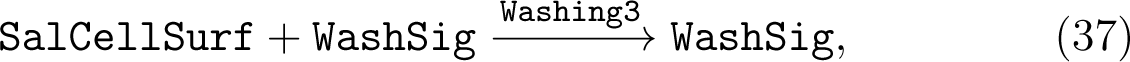

where the places SalMediumStart, SalMedium, and SalCellSurf represent *Salmonella* outside the cell. See Fig. 8 for the graphical representation of the processes (33) to (37).

**Figure 8:**
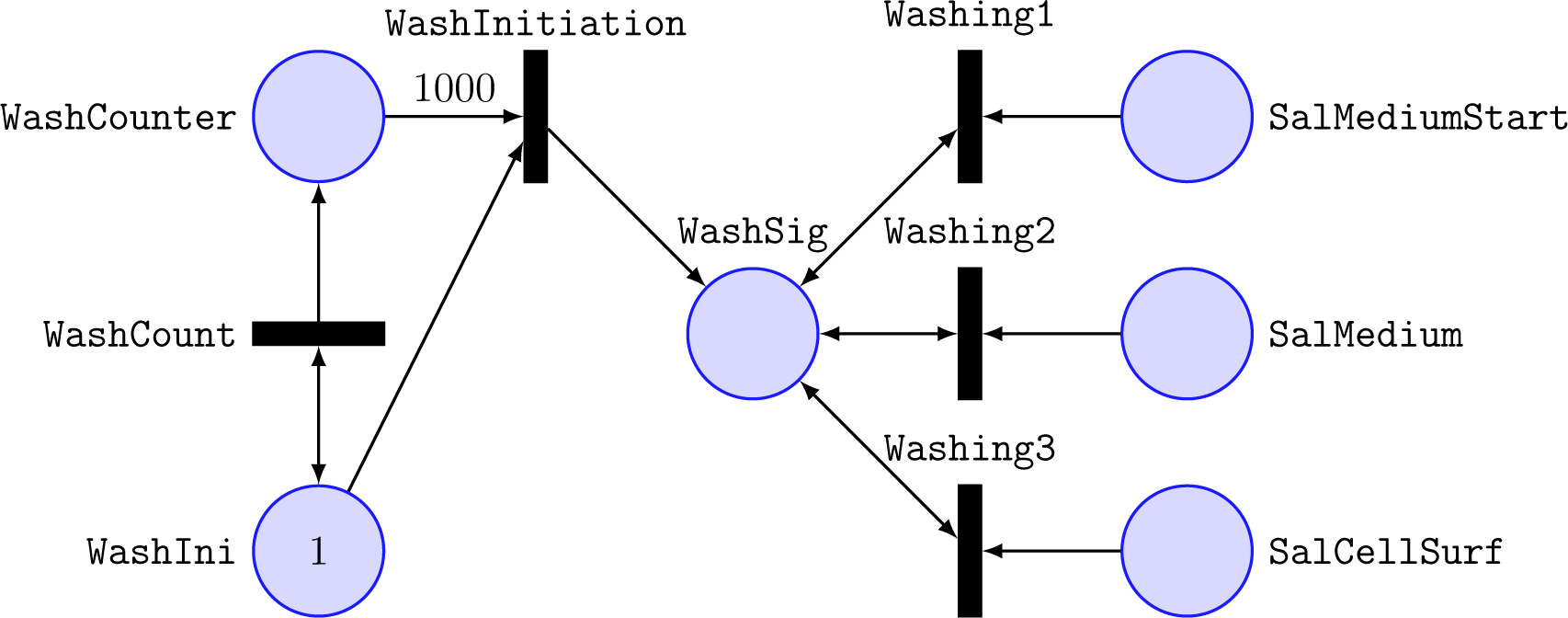
Washing of the epithelial cell to remove the non-invaded bacteria. Circles and rectangles represent the species and processes in (33) to (37), where the associated numbers, 1 and 1000, correspond to the initial marking and the stoichiometric coefficient, respectively.

#### 2.2 Estimation of parameters

Our model parameters, i.e., the rate constants of the processes (1) to (37) denoted as *c*_Transition name_, can be divided into three categories based on the nature of their associated processes. A summary of all parameters is given in Appendix C.

### 2.2.1 Instantaneous processes

In our model, there are processes, which are not concerned with the main stages of the dynamics of the infection process. Therefore, we assume that the occurrence of these processes is instantaneous. This requires a high value to be assigned to their corresponding rate constants, for which we assume a value of 1000 s^−1^. The following rate constants are considered to be in this group: *c*CapBeingReached in (8), *c*AdaptingSigCyt in (15), *c*DividingSalCyt in (17), *c*AdaptingSigVac in (18), *c*DividingSalVac in (20), *c*XenoSigInducing in (22), *c*XenoIniCount in (23), *c*XenoEndCount in (25), *c*DeathCellCyt1 in (27), *c*DeathCellCyt2 in (28), *c*DeathCellVac1 in (29), *c*DeathCellVac2 in (30), *c*Death1 in (31), *c*Death2 in (32), *c*WashInitiation in (34), *c*Washing1 in (35), *c*Washing2 in (36), and *c*Washing3 in (37).

### 2.2.2 Technical processes

The MOI and incubation time are tunable parameters, which model the desired experimental conditions. The first is adjusted through *c*_Adding_ and *c*_Removing_ and the second via *c*_WashCount_. Table 1 gives the values of *c*_Adding_ and *c*_Removing_ for the MOI in the range of 4 to 500, where the generation of the infection medium is considered to be in the timescale of 10^−3^ seconds. Table 2 gives the values of *c*_WashCount_, which adjust the infection time for the range of 5 to 30 minutes.

**Table 1:**
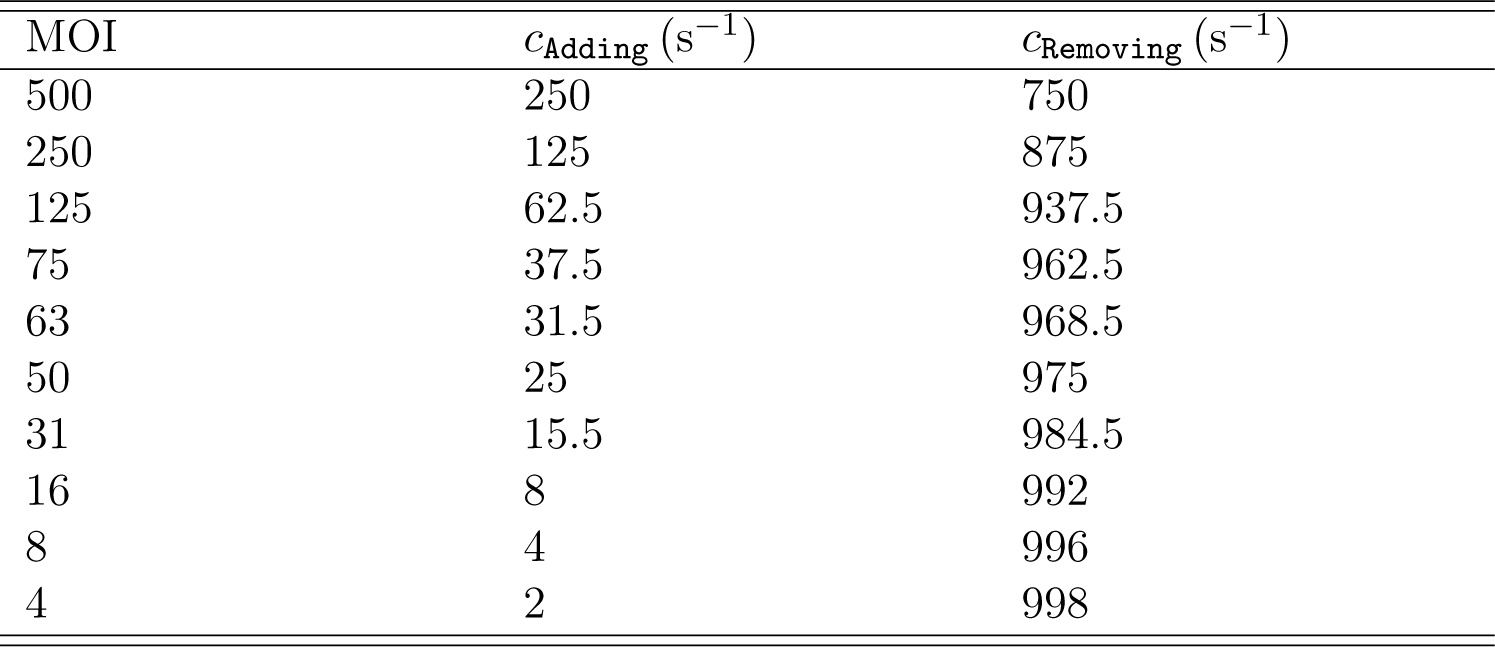
Values of *c*_Adding_ and *c*_Removing_ for various MOIs, given *m*_0_ (MediumGen) = 2000.

**Table 2:** Values of *c*_WashCount_ for various infection times.

| Infection time (min) | $c_{\text{WashCount}}$ ($\text{s}^{-1}$) |
| --- | --- |
| 5 | 3.33 |
| 9 | 1.85 |
| 10 | 1.67 |
| 15 | 1.11 |
| 20 | 0.83 |
| 25 | 0.67 |
| 30 | 0.56 |

### 2.2.3 Dynamical processes

The following are the rate constants, which their associated processes describe the main stages of the infection dynamics: *c*_FirstLanding_ in (3), *c*_TakeOff_ in (4), *c*Landing in (5), *c*Initiation in (6), *c*Joining in (7), *c*StayingVac in (9), *c*EnteringCyt in (10), *c*ProSalVac in (11), *c*ProSalCyt in (12), *c*BeingCytIncapable in (13), *c*BeingCytCapable in (14), *c*AdaptingCyt in (16), *c*AdaptingVac in (19), *c*XenoDeg in (21), *c*_XenoDegInitiation_ in (24), and *c*_XenoDegEnding_ in (26). To estimate these rate constants, we use literature data, and for simulation of the infection dynamics, we use the Gillespie algorithm. Each simulation run corresponds to the fate of an individual cell when exposed to the infection medium characterized by a specific MOI for a given incubation time. The Gillespie algorithm simulates each of the processes in Sec. 2.1 individually and moves forward in time from one process to the next, where its inputs are the initial markings of the places, *m*_0_’s, and the rate constants, *c_i_*’s. This procedure can result in either an infected or uninfected cell. To simulate a population of *n* cells, simulation has to be repeated *n* times. For a detailed discussion of the Gillespie algorithm for Petri net models, we refer to [97]. A common feature of experimental measurements concerns a time period called post-infection (p.i.) time, which represents the time interval between the washing of the epithelial cell and carrying out the measurement. In our model, post-infection time is given by the difference between the simulation and the infection times. For example, for simulation of an experimental measurement of infected cells at 20 minutes post-infection with 10 minutes infection time and an MOI of 50, we consider 30 minutes as the simulation time and the corresponding rate constants of: *c*_Adding_ = 25 s^−1^, *c*_Removing_ = 975 s^−1^, and *c*_WashCount_ = 1.67 s^−1^. Note that, the MOI and the infection time of one individual simulation can differ due to stochastic variations from the average MOI and infection time, listed in Tables 1 and 2, respectively.

#### FirstLanding, TakeOff, and Landing

The median time of near-surface swimming has been observed to be 1.5 seconds [48]. If we assume near-surface swimming as a Poisson process, with an exponentially distributed duration, we obtain *c*_TakeOff_ as: *c*_TakeOff_ = 1*/*1.5 s^−1^ ≈ 0.67 s^−1^.

The concentration of *Salmonella* close to the surface has been measured by fluorescence microscopy, where 2.75×10^−4^ *Salmonella* per µm^2^ have been reported for an MOI of 1.5 [48]. Assuming a HeLa cell with the diameter of 20 µm [98], and thus the area of 314.16 µm^2^, results in 0.0864 *Salmonella* on the average close to the surface of one cell. Under steady-state conditions, the mean number of *Salmonella* on the cell surface of one individual cell is given by

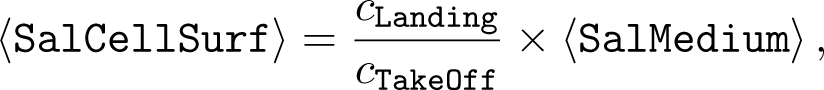

where ⟨· · · ⟩ denotes the average value. The above relation determines *c*_Landing_ = 0.0384 s^−1^, where ⟨SalCellSurf⟩ = 0.0864, *c*_TakeOff_ = 0.67 s^−1^, and ⟨SalMedium⟩ = MOI = 1.5. Since the adaptation of *Salmonella* to the medium, at the beginning of infection, is a slow process, *c*_FirstLanding_ should be smaller than *c*_Landing_. We assume: *c*_FirstLanding_ = *c*_Landing_*/*20 = 0.0019 s^−1^. To check the validity of our estimations, we use them in combination with the parameters estimated in the following, to reproduce the experimental data of [41, 48].

#### Initiation and Joining

In [48], 150 HeLa cells are incubated with *Salmonella* for 9 minutes at various MOIs in the range of 4 to 500. The percentage of cells, possessing at least one membrane ruffle, has been quantified. We have simulated our model 2000 times for the same conditions as in the experiment, that is, 9 minutes infection time and the MOIs in the range of 4 to 500. For the choice of *c*_Initiation_ = 0.0005 s^−1^, simulations, shown in Fig. 9A, reproduce the experimental data of [48] (therein, the left panel of Fig. 10B).

**Figure 9:**
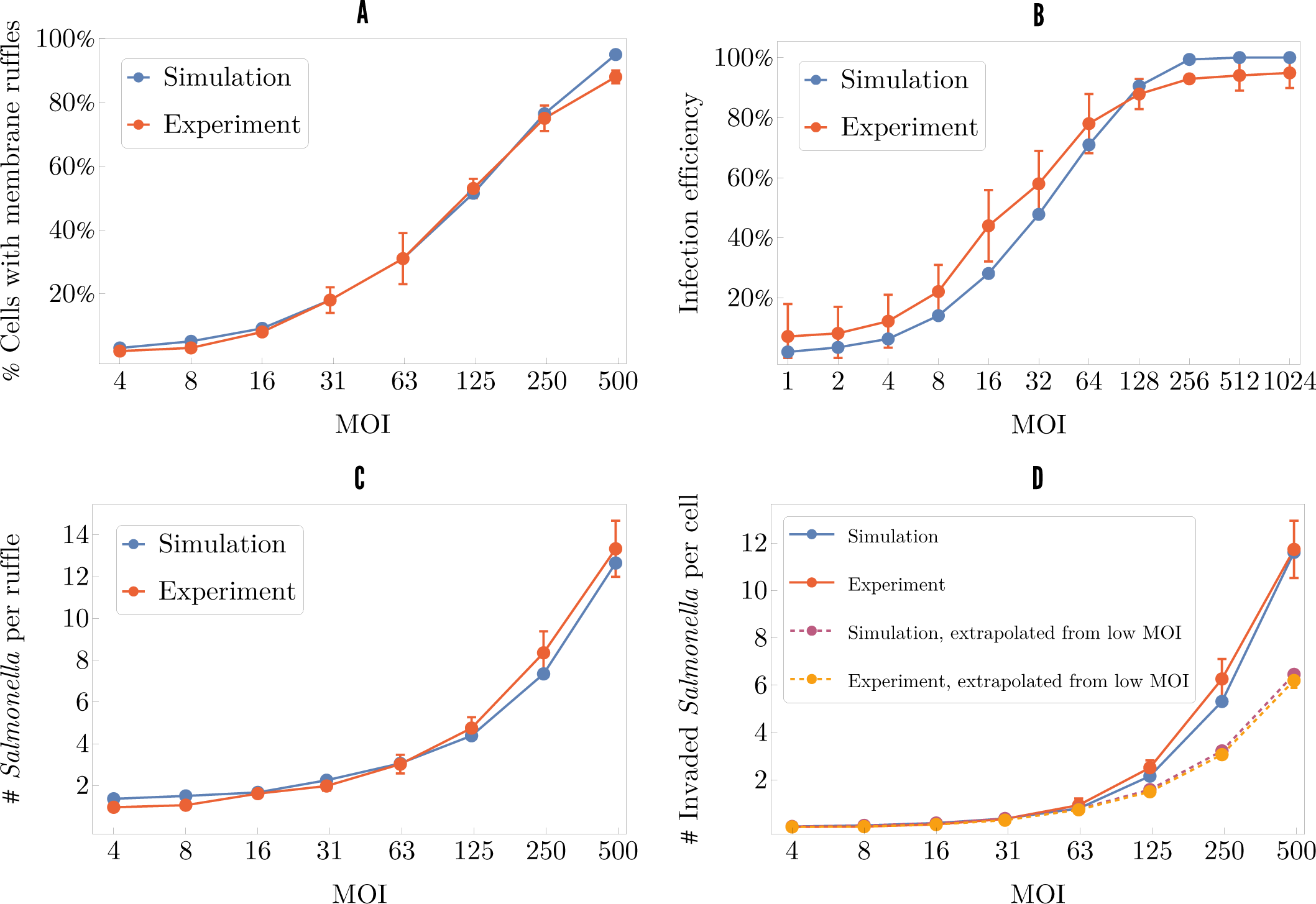
Simulations of the infection dynamics. The experimental data is adopted from [41, 48]. Upper-left panel (A): percentage of cells possessing membrane ruffles (9 minutes infection time). Upper-right panel (B): infection efficiency, i.e., percentage of infected cells (20 minutes infection time). Lower-left panel (C): number of *Salmonella* per ruffle (9 minutes infection time). Lower-right panel (D): number of invaded *Salmonella* per cell. For each MOI, the number of invaded *Salmonella* per cell is calculated by multiplying the fraction of cells, possessing membrane ruffles, with the number of *Salmonella* per ruffle. The dashed lines are extrapolated from low MOIs, assuming a linear increase in invasion efficiency.

**Figure 10:**
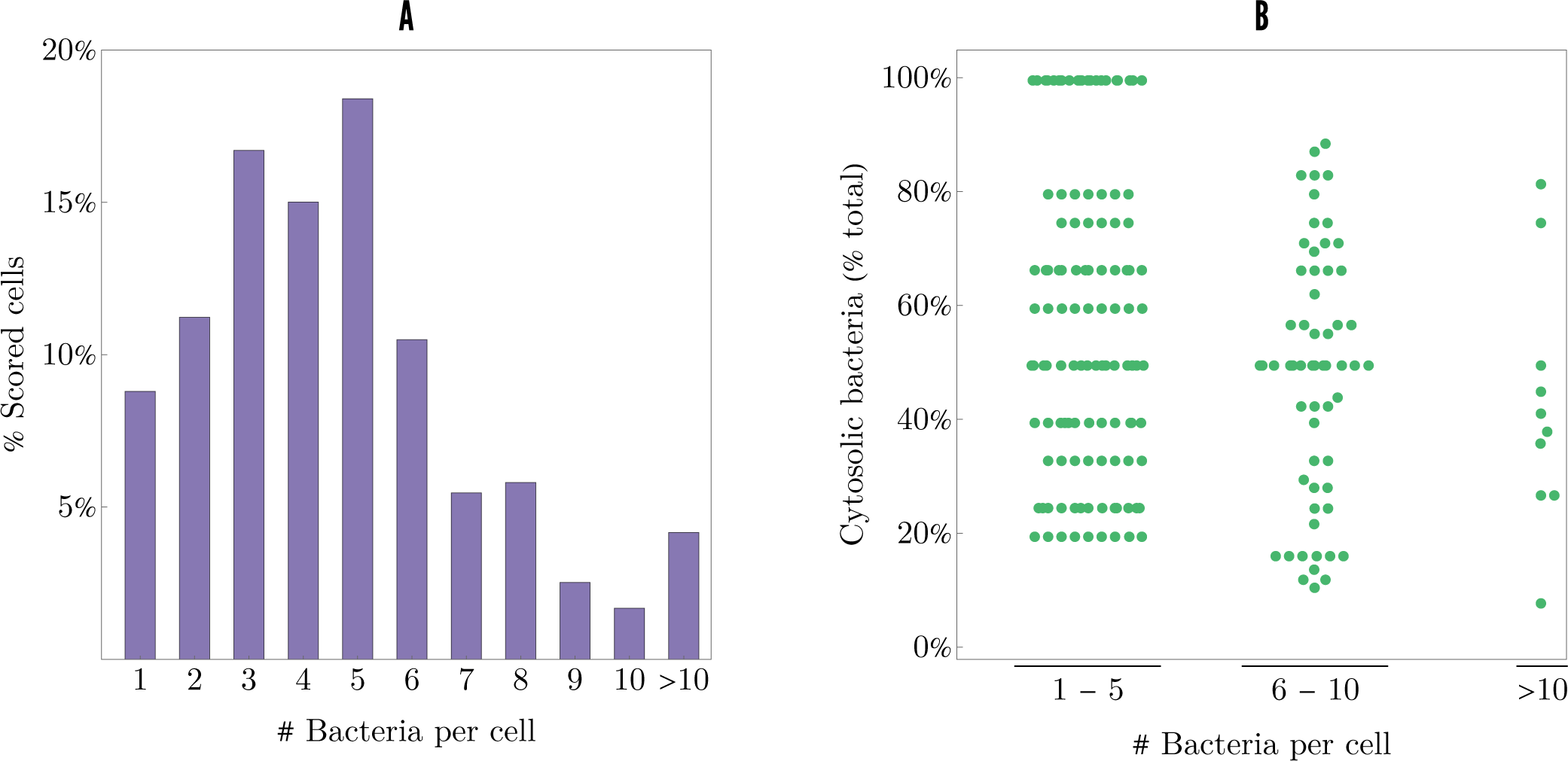
Bacterial load of those cells that contain at least one cytosolic *Salmonella*, i.e., *m* (SalCyt) ≥ 1, for an MOI of 75 and the simulation time of 70 minutes, including 10 minutes infection time. Left panel (A): percentage of cells are scored for each bacterial load, both vacuolar and cytosolic. Right panel (B): fraction of cytosolic *Salmonella* in the total bacterial load for cells, containing 1–5 bacteria, 6–10 bacteria, and *>* 10 bacteria. Each dot depicts one simulation, that is, one infected cell. These results are in agreement with the experimental data of [55] and confirm the hypothesis that there is no correlation between vacuolar lysis and bacterial load.

Formation of ruffles requires the irreversible binding of at least one bacterium [40]. The percentage of cells, possessing ruffles, can be considered as the proportion of infected cells. Figure 9B shows the infection efficiency defined as the fraction of the infected cells for the incubation time of 20 minutes and the MOI in the range of 1 to 1024. The simulated infection efficiency is in agreement with [41] (therein, Fig. 1E).

To model the significant cooperative effect of *Salmonella* invasion [48, 57], the probability of bacteria, joining preexisting ruffles, has to be chosen higher than the probability of the formation of new ruffles, i.e., *c*_Joining_ *> c*_Initiation_. We adjust *c*_Joining_ to reproduce the middle panel of Fig. 10B in [48], where the number of *Salmonella* per ruffle, above (outside) and under (inside) the cell surface, has been quantified four times in 25 HeLa cells. Following the experimental conditions, we have simulated our model with an infection time of 9 minutes at various MOIs in the range of 4 to 500. For each MOI, the number of bacteria per membrane ruffle is enumerated in simulations with at least one ruffle, i.e., *m* (NrRuffle) ≥ 1. For *c*_Joining_ = 0.005 s^−1^, which is 10 times higher than *c*_Initiation_ = 0.0005 s^−1^, the model reproduces the experimental data of [48], see Fig. 9C. In our simulations, in contrast to [48], we have not distinguished between the bacteria above and under the cell surface. To compare our results with the experimental data, we have summed up the number of bacteria above and under the cell surface and have called the resulting quantity number of *Salmonella* per ruffle. Figure 9C shows that number of *Salmonella* per ruffle increases with enhancing MOI. For the MOI in the range of 63 to 125, there are three to five bacteria located in one ruffle. For a high MOI, e.g., 500, the number of bacteria clustering in a ruffle reaches 13.

Similar to [48] (therein, the right panel of Fig. 10B), we have multiplied the fraction of cells, possessing membrane ruffles, with the number of *Salmonella* per ruffle, and obtained the number of invaded *Salmonella* per cell. Figure 9D shows the number of invaded *Salmonella* per cell for the MOI in the range of 4 to 500. By the assumption of no cooperative behavior of the bacterial invasion, a linear increase of invasion efficiency would be expected with increasing MOI [48]. By taking this into account, we have extrapolated from low to high MOIs—dashed lines in the plot—where the extrapolated line is much lower than the simulation values, which indicates the cooperative invasion of *Salmonella*.

It can be assumed that the area of the ruffled membrane, containing multiple bacteria, is considerably larger than the size of a ruffle with one or only a few bacteria. An increase in the ruffle size, which implies an expansion of the size of the obstacle on the cell surface, may enhance the cooperative effect of the ruffle-joining by further bacteria. Since our model simulations are in quantitative agreement with the experimental data, we have not taken into account the relation between the geometric size of a ruffle and the act of ruffle-joining.

#### StayingVac and EnteringCyt

The proportion of vacuolar or cytosolic *Salmonella* in cytosolic-capable cells is determined by *c*_StayingVac_ and *c*_EnteringCyt_, respectively. We choose the values of these two parameters to reproduce the experimental data presented in Fig. 2 of [55]. Therein, HeLa cells are infected by *Salmonella* at the MOI in the range of 50 to 100, and the bacterial load and fraction of cytosolic *Salmonella* in cells that contain at least one cytosolic bacterium have been quantified at one-hour post-infection. In [55], the bacterial load is defined as the total number of bacteria, including both vacuolar and cytosolic per cell. Note that, the experimental data of [55] does not reflect the situation immediately after the infection. At one-hour post-infection, the effects of proliferation and xenophagy become significant. According to the experimental conditions, we have simulated our model at an average MOI of 75 for 70 minutes (including 10 minutes of infection time). We have performed 3000 simulation runs, and each simulation run represents the infection of one individual cell, which have resulted in *>* 150 cells with cytosolic *Salmonella*, i.e., *>* 150 cells with *m* (SalCyt) ≥ 1. For these cytosolic-capable cells, the total number of internalized bacteria and the fraction of cytosolic *Salmonella* in the total bacterial load are shown in Fig. 10. Except for the stochastic variations, the simulation results are in agreement with the experimental data for *c*_StayingVac_ = 0.006 s^−1^ and *c*_EnteringCyt_ = 0.004 s^−1^ (Fig. 2 of [55]). Similar to the distribution measured experimentally, the simulated bacterial load in cells with cytosolic *Salmonella* is broadly distributed, ranging from 1 to more than 10 bacteria per cell, see Fig. 10A. The fraction of cytosolic *Salmonella* in cells with high bacterial loads, that is, 6–10 bacteria and *>* 10 bacteria per cell, is mostly lower than the fraction of vacuolar *Salmonella*, see Fig. 10B. If a correlation between the SCV damage and the bacterial load exists, we would expect a higher number of cytosolic bacteria in the cells with a high bacterial load. The simulations confirm the hypothesis proposed in [55] that no correlation exists between the vacuolar lysis and the bacterial load of cells.

#### ProSalVac and ProSalCyt

For the adjustment of the proliferation rate of vacuolar and cytosolic *Salmonella*, we need to determine *c*_ProSalVac_ and *c*_ProSalCyt_. The proliferation inside the cytosol has been demonstrated to be much faster than inside the SCV, due to ideal living conditions, such as a nutrient-rich environment, sufficient space, and neutral pH [27, 28, 34, 39, 55, 58]. The term hyper-replication describes this phenotype of fast-replicating *Salmonella*, which results in more than 50–100 bacteria per cell [39, 55]. In human colonic epithelial cells, the doubling time of ∼ 20 minutes for cytosolic *Salmonella* has been observed [39]. In HeLa cells, the growth of cytosolic *Salmonella* is investigated in [46], where to distinguish between vacuolar and cytosolic bacteria, fluorescent dextran is used, which is a marker of the endocytic pathway that accumulates within the SCV [34]. To reproduce the experimental data presented in Fig. 4A of [46], we choose a slightly slower doubling time of 40 minutes for cytosolic *Salmonella* in HeLa cells, which corresponds to the proliferation rate of 0.000417 *Salmonella* per second and determines *c*_ProSalCyt_ as: *c*_ProSalCyt_ = 0.000417*/*Max (DiviSalCyt) s^−1^ = 0.000000595 s^−1^, where Max (DiviSalCyt) = 700. The simulated growth of *Salmonella* in the cytosol is depicted in Fig. 11A, which shows the fold change versus 2.5 hours postinfection of cytosolic *Salmonella* for ten simulation runs, i.e., ten infected cells, with *m* (SalCyt) ≥ 1. The simulated cytosolic growth resembles the experimental growth of dextran-negative (cytosolic) *Salmonella* in Fig. 4A of [46]. Similar to the experimentally observed cytosolic growth, the late onset of proliferation between four and six hours post-infection is predicted by our model. In the simulations, the late onset is caused by the xenophagic degradation of cytosolic *Salmonella* in the first hours of the infection and not by, for example, the long lag phase of cytosolic bacteria, which is in agreement with the detection of cytosolic bacterial growth as early as four hours post-infection in NDP52-depleted cells, see [38].

**Figure 11:**
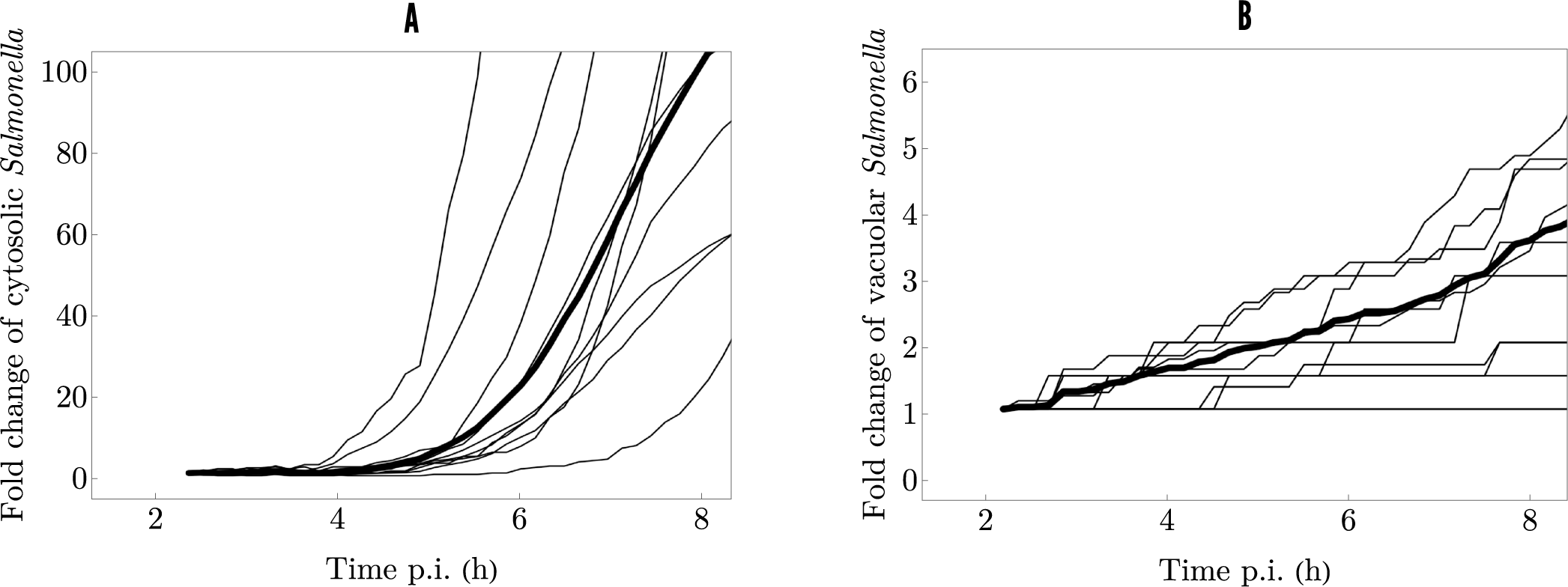
Simulated growth of cytosolic and vacuolar *Salmonella*. The fold change versus 2.5 hours post-infection of (left panel) cytosolic *Salmonella* and (right panel) vacuolar *Salmonella* are quantified at 10 minutes intervals for the indicated time points with 10 minutes infection time and an MOI of 50. The mean fold change is depicted by a thick line in each case. The simulated growth of *Salmonella* resembles the experimental observation reported in [46].

In human colonic epithelial cells, the average doubling time of *Salmonella* inside the SCV has been measured to be ≥ 95 minutes [39]. Consequently, the proliferation inside the SCV has to be much slower than inside the cytosol. A three-fold increase for vacuolar *Salmonella* measured in [46] from 4–5 to 9 hours post-infection, which corresponds to the doubling time of approximately 4 hours and 30 minutes for vacuolar bacteria, results in the proliferation rate of 0.0000617 *Salmonella* per second. To achieve the maximum doubling time of 4 hours and 30 minutes for a bacterium, *c*_ProSalVac_ is set to: *c*_ProSalVac_ = 0.0000617*/*Max (DiviSalVac) s^−1^ = 0.000000411 s^−1^, where Max (DiviSalVac) = 150. The simulated growth of vacuolar *Salmonella* is shown in Fig. 11B, where for ten simulation runs, with *m*(SalVac) ≥ 1, the fold change of vacuolar *Salmonella* is quantified. Figure 11B resembles the experimental growth of dextran-positive (vacuolar) *Salmonella* depicted in Fig. 4B of [46].

#### BeingCytIncapable and BeingCytCapable

The chance of a cell to become cytosolic-capable or cytosolic-incapable is determined by the ratio of *c*_BeingCytCapable_ and *c*_BeingCytIncapable_. The fraction of cells with hyper-replicating *Salmonella*, cells containing cytosolic *Salmonella*, has been determined to be around 8% for various MOIs in the range of 25 to 800 [95]. In another study, ∼ 10% of cells are detected to contain ubiquitylated bacteria at two and four hours post-infection [38]. Ubiquitin has been demonstrated to serve as a marker for the presence of *Salmonella* in the cytosol [30]. It is reported that approximately 9% of HeLa cells harbor hyper-replicating *Salmonella* at 8 hours post-infection [55]. The study of [46] demonstrates that at 6 hours post-infection ∼ 15% of infected cells contain cytosolic *Salmonella* stained with anti-LPS antibody. Therein, it is also reported that at 8 hours post-infection approximately 10% of cells contain hyper-replicating *Salmonella*.

Xenophagy has been shown to restrict the proliferation of *Salmonella* in the cytosol [38, 42]. Therefore, the fraction of 8% to 15% of cells, containing cytosolic bacteria, is not necessarily the fraction of cytosolic-capable cells. The autophagy receptor NDP52 is depleted and the fraction of HeLa cells containing ubiquitin-coated *Salmonella* has been measured for two and four hours post-infection in [38]. At two hours post-infection ∼ 23% of cells contain cytosolic *Salmonella* and ∼ 45% at four hours post-infection. Based on these experimental findings, we assume that around 35% of the cells are cytosolic-capable. To generate 35% cytosolic-capable and 65% cytosolic-incapable cells, we consider: *c*_BeingCytCapable_ = 350 s^−1^ and *c*_BeingCytIncapable_ = 650 s^−1^.

#### XenoDeg, XenoDegInitiation, and XenoDegEnding

To specify in which period of time xenophagy is active, i.e., *m* (XenoSig) = 1, we need to adjust *c*_XenoDegInitiation_ and *c*_XenoDegEnding_. It is suggested that xenophagy is initiated by amino-acid starvation, which in turn is provoked by membrane damage and is only active in the early stages of the infection process [47,50]. Furthermore, at one-hour post-infection, approximately 20% of total intracellular bacteria are already associated with LC3 proteins [32]. We choose *c*_XenoDegInitiation_ = 0.56 s^−1^ to activate xenophagy, i.e., *m* (XenoSig) = 1, after ∼ 30 minutes, when the first bacterium reaches the cytosol or when damage to the SCV is first initiated. At three to four hours post-infection, xenophagy has been reported to be prevented by normalization of cytosolic amino-acid levels and reactivation of mTOR [47, 50]. Choosing the value of *c*_XenoDegEnding_ = 0.069 s^−1^ leads to the deactivation of xenophagy, i.e., *m* (XenoSig) = 0, after ∼ 4 hours when the first bacterium reaches the cytosol.

The efficiency of xenophagic degradation is determined through *c*_XenoDeg_. We estimate its value by reproducing the experimental data of [46,55], where 10 ± 4 (mean ± SD) % and 9.2 ± 3.2% of infected HeLa cells are reported to contain hyper-replicating *Salmonella* at 8 hours post-infection. We can predict the fraction of 8.7% of cells with cytosolic, hyper-replicating *Salmonella* at 8 hours post-infection for *c*_XenoDeg_ = 0.00037 s^−1^, see Sec. 3 and Fig. 13.

**Figure 12:**
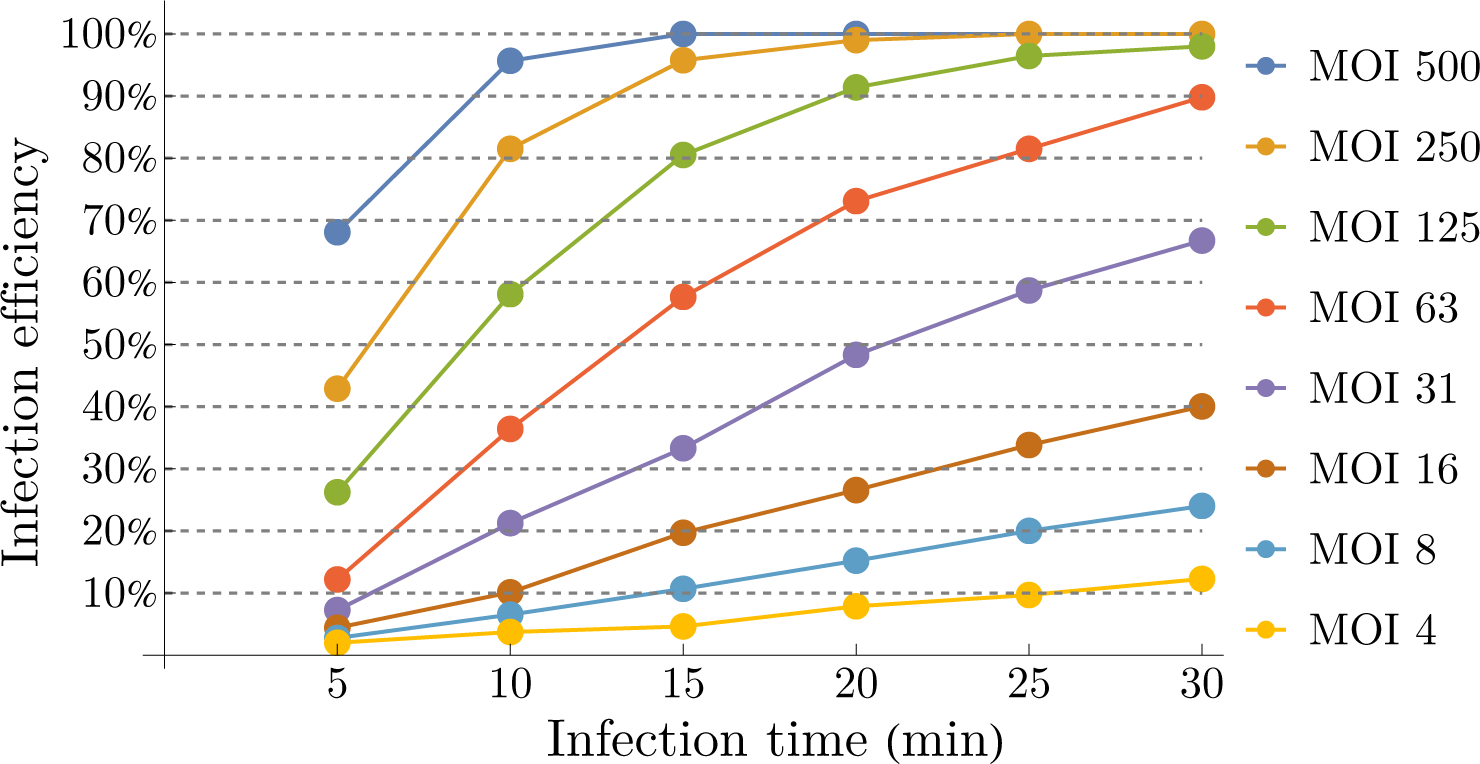
Predicted infection efficiency of infected HeLa cells using our model.

**Figure 13:**
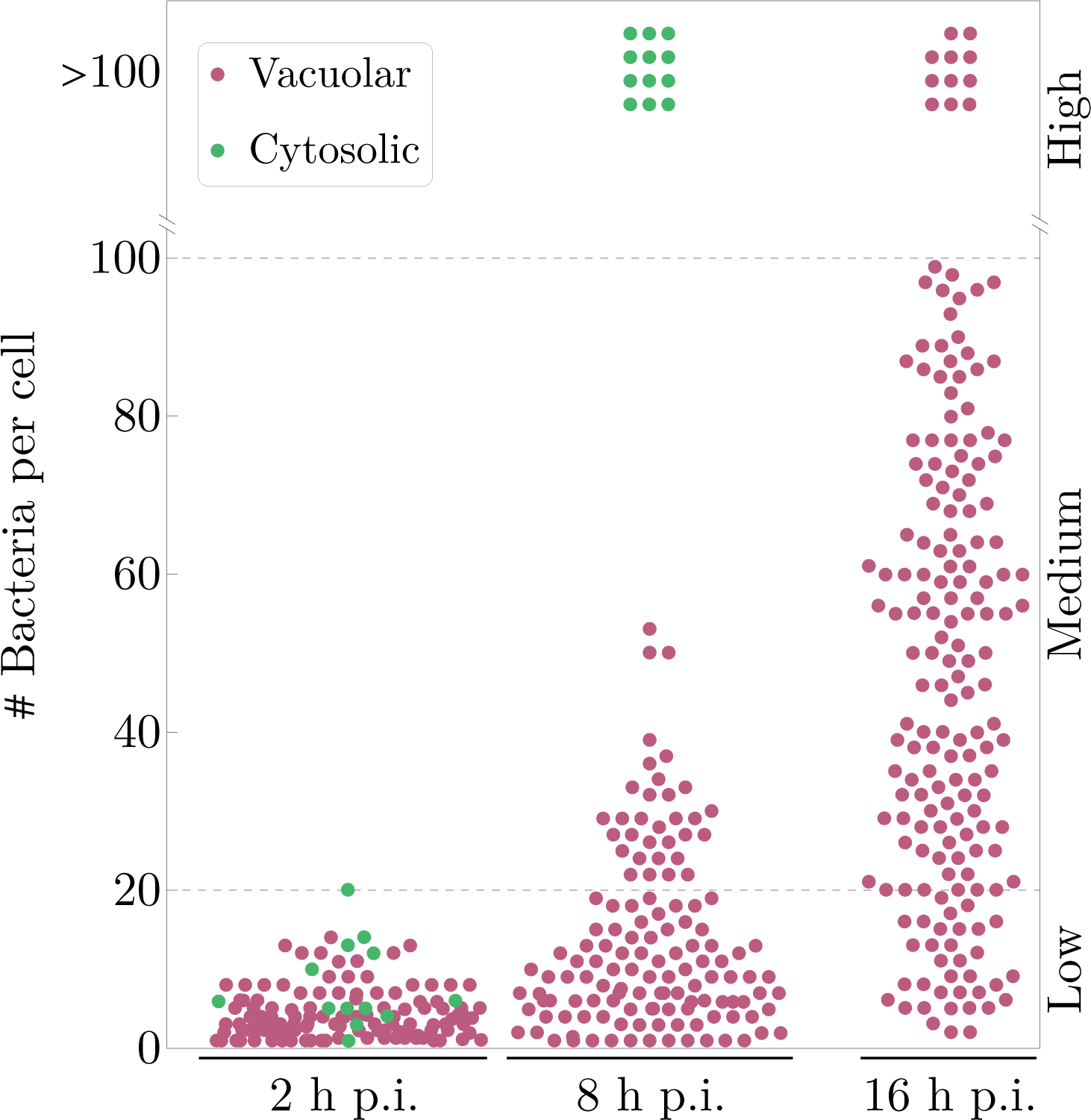
Prediction of the intracellular bacterial composition. The number of bacteria is counted per infected cell at 2, 8, and 16 hours post-infection with *>* 100 cells, 10 minutes infection time, and an MOI of 50. Each dot represents one infected cell in maroon for cells, which host exclusively vacuolar bacteria, and in green for those that also contain cytosolic bacteria.

#### AdaptingCyt and AdaptingVac

By determining *c*_AdaptingCyt_ and *c*_AdaptingVac_, the time period of the lag phases of cytosolic and vacuolar *Salmonella* can be adjusted. However, there are no experimental data available concerning the lag phase of *Salmonella* inside HeLa cells. It has been observed that bacterial growth in the cytosol is initiated around four hours post-infection which is, however, mainly caused by the xenophagic degradation in the first hours of infection and not by a long lag phase. To get the highest achievable doubling time with *m* (DiviSalCyt) = 700 after ∼ 70 minutes when the first bacterium reaches the cytosol, we set: *c*_AdaptingCyt_ = 0.167 s^−1^. Similarly, in the case of vacuolar bacteria with Max (DiviSalVac) = 150, we set: *c*_AdaptingVac_ = 0.036_s_−1.

## 3 Predictions

Here, we present two predictions of our model, the effects of the incubation time and the MOI on the infection efficiency of epithelial cells and the bacterial load of the infected cells at different post-infection times.

### Infection efficiency

In Fig. 12, we have depicted the prediction of our model for the infection efficiency of infected HeLa cells, for the infection time ranging from 5 to 30 minutes and the MOI from 4 to 500. Figure 12 shows that the infection efficiency enhances by increasing the MOI and infection time, as expected. For example, for the high MOI of 500, nearly 100% of the cells are predicted to be infected from the infection time of 15 minutes onward. For a low MOI, e.g., an MOI of 4, even for a high infection time of 30 minutes, only around 12% of the cells are infected. Depending on the context and the biological questions, the ideal MOI and infection time have to be selected for the experiments. For example, a low MOI can be experimentally motivated by the avoidance of multiple infection events of an individual cell. Our model can provide useful information in this regard and can be used to predetermine the optimal experimental setting in infectious cell cultures.

### Composition of the bacterial load

To investigate the composition and distribution of the intracellular bacteria per cell over time, we have exploited our model to study the number of internalized bacteria for over 100 infected cells at different post-infection times, see Fig. 13. Each dot represents the total number of *Salmonella* in one cell. The simulation results are in agreement with the experimental data shown in Fig. 1B of [46] and in Fig. 1 of [55], where in the experiments, the intracellular *Salmonella* are enumerated by fluorescence microscopy and no distinction has been made between vacuolar and cytosolic *Salmonella*. In our simulation, we can differentiate between these two. A maroon dot in Fig. 13 depicts the number of bacteria of one individual cell with exclusively *Salmonella* in the SCV, and a green dot represents the total number of bacteria per cell, including cytosolic *Salmonella*.

At 2 hours post-infection, all cells contain less than 20 bacteria, independent of their locations in the SCV or cytosol. At 8 hours post-infection, cells have separated into two groups. The first group comprises the cells, hosting exclusively vacuolar bacteria, where most of them still contain less than 20 bacteria (the maximum corresponds to 53 bacteria in one cell). By contrast, in the second group, all cells contain more than 100 bacteria. In [46, 55], it is suggested that this group represents the hyper-replicating cytosolic *Salmonella*. Our model confirms this hypothesis. Note that these cells can also contain vacuolar *Salmonella*. A proportion of 8.7% of the cells belongs to the second group containing cytosolic *Salmonella*, which is in agreement with the fraction of the hyper-replicating bacteria measured in the experiments: 10 ± 4 (mean ± SD) % in [46] and 9.2 ± 3.2% in [55]. At 16 hours post-infection, the number of bacteria per cell is broadly distributed, ranging from 2 to more than 100 *Salmonella* per cell. At this stage of infection, all cells are predicted to host exclusively vacuolar bacteria, that is, all cells that used to contain cytosolic *Salmonella* have already died at this late phase of infection. Here, cells with more than 100 *Salmonella* per cell are predicted to be the cells, which contain bacteria enclosed in their SCVs.

The term hyper-replication, concerning the cells with more than 50–100 bacteria per cell, has been proposed to describe the fast-replicating *Salmonella* inside the cytosol of these cells [39, 55]. Our model demonstrates that the slow-growing vacuolar bacteria can also reach more than 50–100 bacteria per cell at the later time point of infection. According to the definition of hyper-replication in [39, 55], these bacteria would be classified as such and thereby misidentified as cytosolic *Salmonella*. Our simulation demonstrates that the term hyper-replication based on the number of bacteria per HeLa cell for describing cytosolic *Salmonella* is only valid at 8 hours post-infection, but not at 2 and 16 hours.

## 4 Summary and outlook

Gram-negative bacteria of the *Salmonella* genus can affect almost all major organs, such as the liver, called *Salmonella* hepatitis [99, 100], spleen [100, 101], gall bladder [102], and the central nervous system, *Salmonella* meningitis [103]. Additionally, *Salmonella* has also been frequently used as a model organism to study the mechanisms involved in bacterial infections. Due to the high variability and heterogeneity of the vacuolar and cytosolic *Salmonella*, non-deterministic models are suitable for capturing the stochastic nature of bacterial infection at a single-cell level.

In this paper, by assuming the infection of epithelial cells by *Salmonella* as a discrete-state, continuous-time Markov process and applying the Gillespie algorithm, we construct the first stochastic model for the time evolution of *Salmonella* infection at its different stages of development at a cellular level. To estimate the parameters of our model, we used human epithelial cell line (HeLa cells) data reported in the literature. Besides reproducing the experimental findings by assigning biologically sensible values to the model parameters, we predict the percentage of infected cells, depending on the two factors of infection time and MOI, and shed light on the dynamics of bacterial replication in the SCV and cytosol. Our model simulations demonstrate that there is no correlation between the vacuolar lysis and the bacterial load of cells. At the later time points of infection, the proliferation of *Salmonella* in the SCV resembles hyper-replicating cytosolic bacteria. This finding challenges the common belief that associates the phenomenon of hyper-replication with cytosolic bacteria.

The question arises whether *in vitro* studies of *Salmonella* infection, using HeLa cells as a model system, can elucidate and reveal the processes that occur under *in vivo* conditions. Most of the processes considered here have been observed to occur *in vivo* in similar ways. For example, the cooperative invasion has been detected in polarized epithelial cells [57] and also in the intestinal epithelium of guinea pigs [6, 7] though not in the murine gut [63]. The aggregation of multiple bacteria on a single ruffle has been observed in cultured intestinal epithelial cells [22]. The near-surface swimming has been shown by *in vivo* intravital microscopy in the cecum of infected mice [44]. The study of polarized human intestinal epithelial cells in [39] has demonstrated that *Salmonella* in the cytosol proliferate at higher rates than in the SCV. The extrusion of these cells from the monolayer is associated with pyroptosis and results in the dissemination of bacteria into neighboring cells.

Future experiments can help improve the estimation of the parameters of our model. Quantitative single-cell experiments in combination with live-cell imaging and computational image analysis may monitor when a bacterium invades, proliferates, and is eliminated by xenophagy. The use of markers, concerning the localization of intracellular bacterium, would be essential, e.g., lysosome-associated membrane protein 1 (LAMP1) [10] or dextran to visualize the SCV [34]. Furthermore, to gain a better understanding of how xenophagy targets *Salmonella* inside the cytosol, it is necessary to shift from population-based studies towards single-cell or even single-bacterium analysis [104].

The approach proposed in this paper can be modified to take into account new features or mechanisms, concerning the invasion of epithelial cells by *Salmonella*, or to model the infection dynamics of other pathogens. For example, recently it has been suggested that the cooperative invasion through expansive ruffles might only be an *in vitro* process, as it is not detected in the intact mouse gut [63]. To modify our model accordingly, the module, concerning the cooperative invasion, can be simply turned off, which is achieved by setting *c*_Joining_ = 0. Similarly, in the case of other pathogens, such as *Shigella flexneri*, which show a similar behavior as *Salmonella* in invading host cells [37], only the estimation of parameters needs to be adjusted.

## Acknowledgments

We acknowledge funding from the consortia ACLF-I (Acute-on-Chronic Liver Failure - Initiative) and ENABLE (Unraveling mechanisms driving cellular homeostasis, inflammation, and infection to enable new approaches in translational medicine) (Hessian Ministry for Science and the Arts). We would like to thank Goethe University Frankfurt and the Hessian Ministry of Higher Education, Research, and the Arts, for providing financial and infrastructural support.

## Appendix A Processes

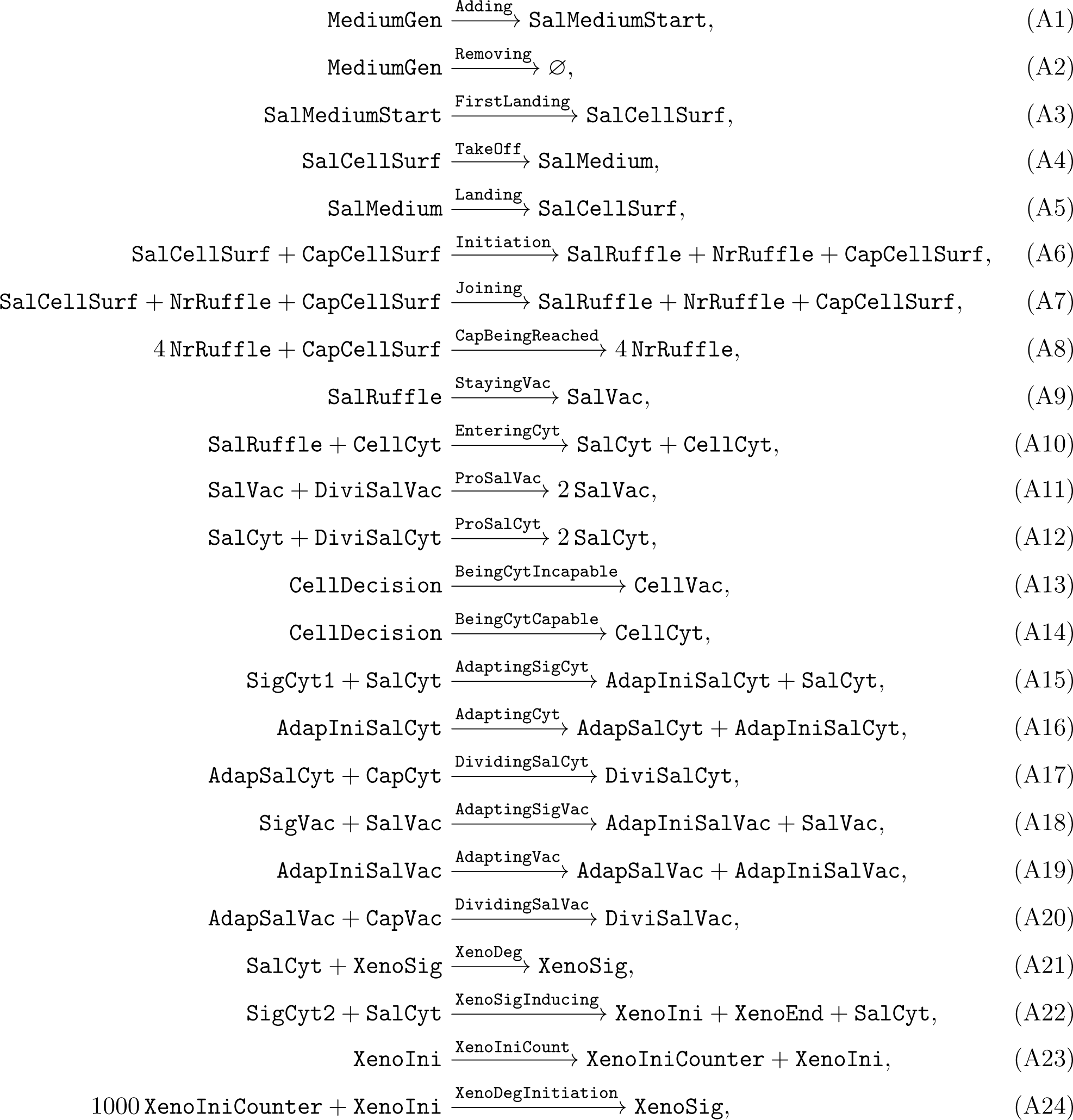
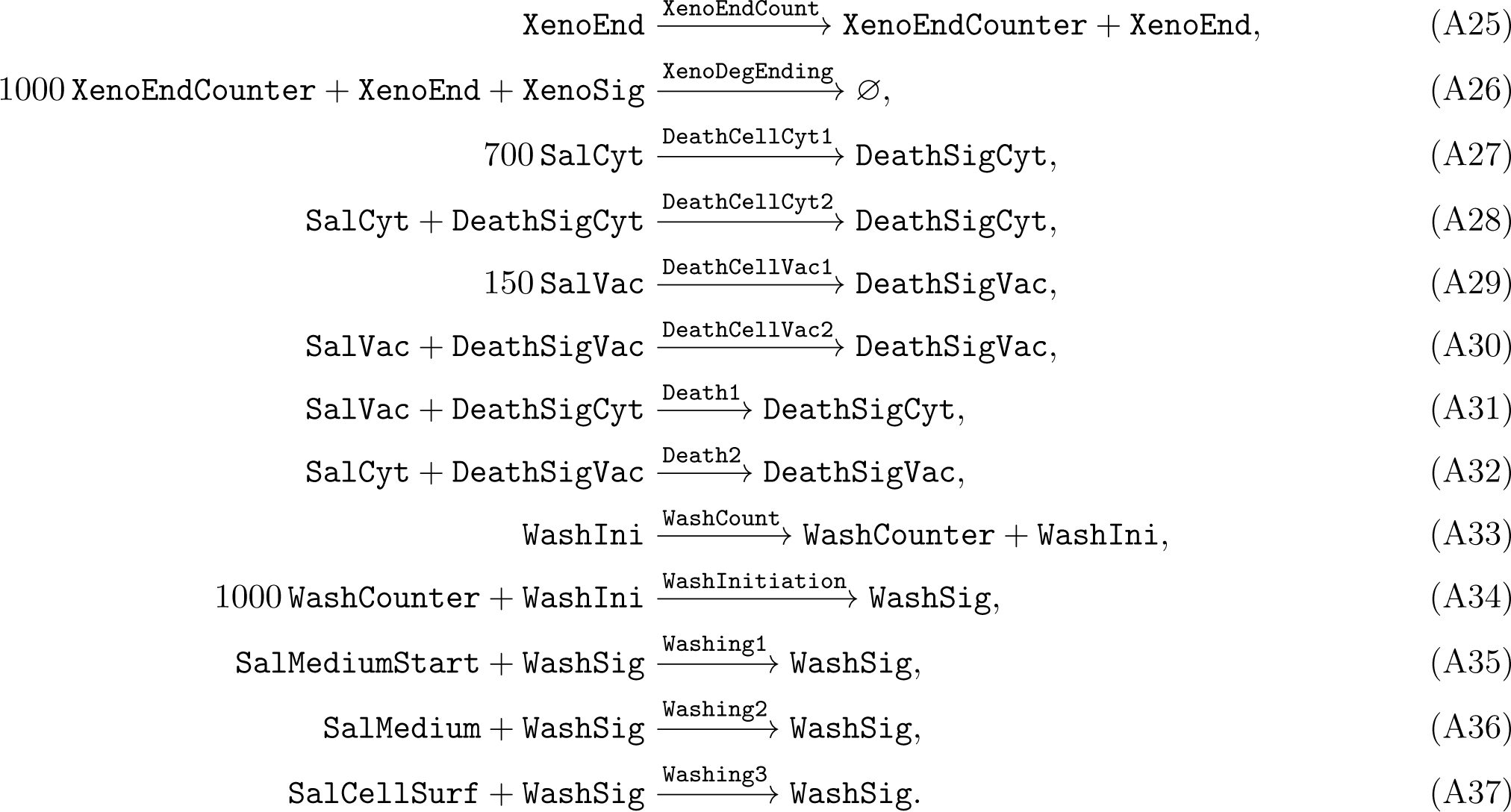

## Appendix B Places of the Petri nets with their initial markings

| Place | Description | Initial marking ($m_0$) |
| --- | --- | --- |
| MediumGen | Maximum possible number of bacteria to infect an individual cell | 2000 |
| SalMediumStart | Actual number of *Salmonella* in the medium to infect an individual cell | 0 |
| SalCellSurf | Number of bacteria on the cell surface | 0 |
| SalMedium | Number of the adapted bacteria to the medium | 0 |
| CapCellSurf | Signal for the capability of the cell surface to accommodate additional bacteria | 1 |
| SalRuffle | Total number of bacteria located in ruffles | 0 |
| NrRuffle | Counter for the number of ruffles on the cell surface | 0 |
| SalVac | Total number of *Salmonella* in the SCV | 0 |
| CellCyt | Cytosolic-capable cell | 0 |
| SalCyt | Total number of *Salmonella* in the cytosol | 0 |
| DiviSalVac | Divisibility of vacuolar *Salmonella* | 0 |
| DiviSalCyt | Divisibility of cytosolic *Salmonella* | 0 |
| CellDecision | Cell before making a choice to become cytosolic-capable or cytosolic-incapable | 1 |
| CellVac | Cytosolic-incapable cell | 0 |
| SigCyt1 | Signal for the presence of *Salmonella* in the cytosol | 1 |
| AdapIniSalCyt | Signal for the initiation of the adaptation process of *Salmonella* to the cytosol | 0 |
| AdapSalCyt | Adjustment to the cytosol | 0 |
| CapCyt | Capacity of the cytosol | 700 |
| SigVac | Signal for the presence of *Salmonella* in the vacuole | 1 |
| AdapIniSalVac | Signal for the initiation of the adaptation process of *Salmonella* to the vacuole | 0 |
| AdapSalVac | Adjustment to the vacuole | 0 |
| CapVac | Capacity of the SCV | 150 |
| XenoSig | Signal for xenophagic degradation | 0 |
| SigCyt2 | Signal for the presence of *Salmonella* in the cytosol | 1 |
| XenoIni | Signal for the initiation of xenophagy | 0 |
| XenoEnd | Signal for the termination of xenophagy | 0 |
| XenoIniCounter | Time counter for the activation of xenophagy | 0 |
| XenoEndCounter | Time counter for the deactivation of xenophagy | 0 |
| DeathSigCyt | Signal that cell undergoes death due to a high number of cytosolic *Salmonella* | 0 |
| DeathSigVac | Signal that cell undergoes death due to a high number of vacuolar *Salmonella* | 0 |
| WashIni | Signal for the one-time initiation of the washing process | 1 |
| WashCounter | Time counter of the washing process | 0 |
| WashSig | Signal for the onset of the washing process | 0 |

## Appendix C Transitions of the Petri nets with their rate constants

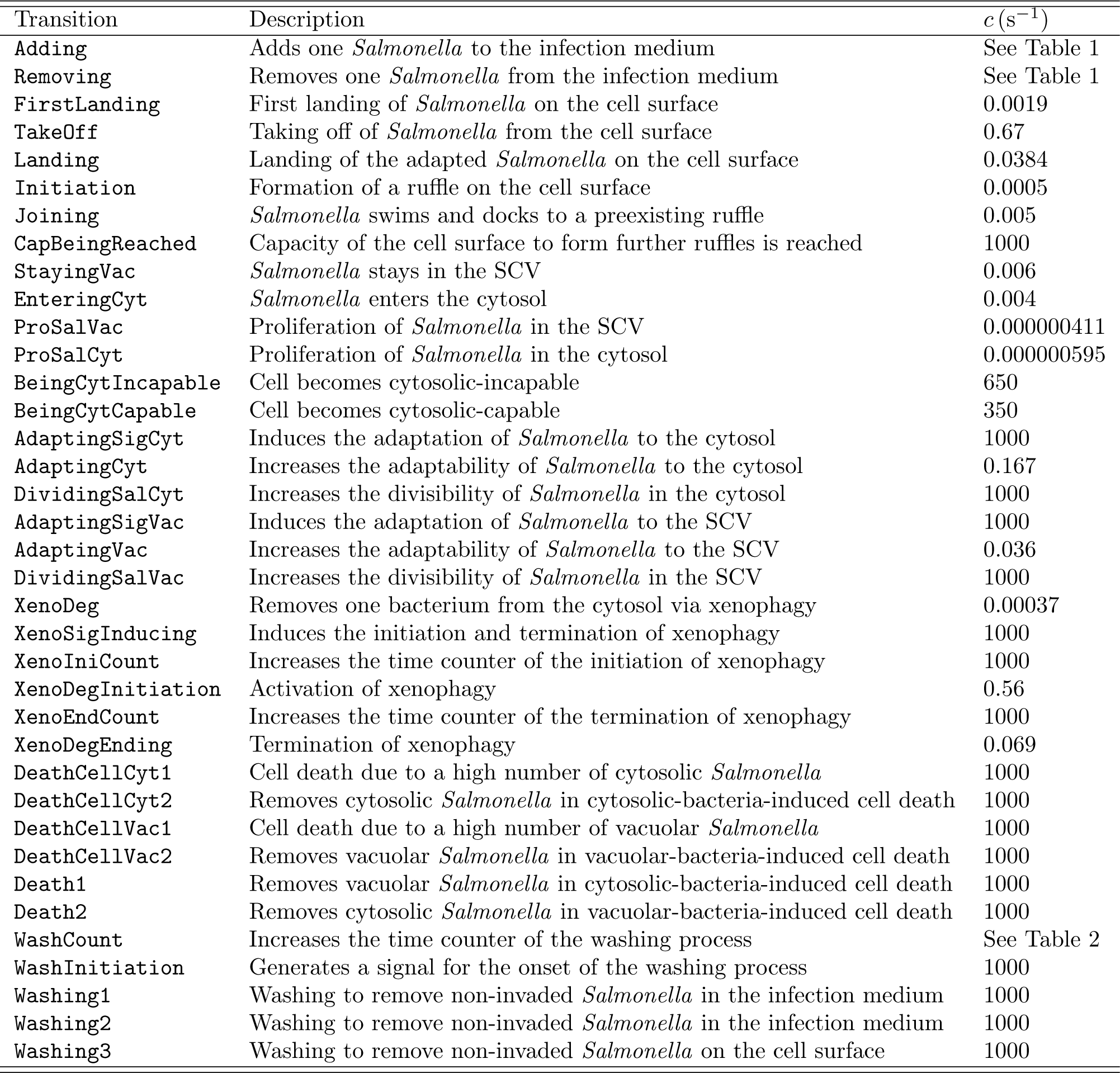

| Transition | Description | $c (\text{s}^{-1})$ |
| --- | --- | --- |
| Adding | Adds one *Salmonella* to the infection medium | See Table 1 |
| Removing | Removes one *Salmonella* from the infection medium | See Table 1 |
| FirstLanding | First landing of *Salmonella* on the cell surface | 0.0019 |
| TakeOff | Taking off of *Salmonella* from the cell surface | 0.67 |
| Landing | Landing of the adapted *Salmonella* on the cell surface | 0.0384 |
| Initiation | Formation of a ruffle on the cell surface | 0.0005 |
| Joining | *Salmonella* swims and docks to a preexisting ruffle | 0.005 |
| CapBeingReached | Capacity of the cell surface to form further ruffles is reached | 1000 |
| StayingVac | *Salmonella* stays in the SCV | 0.006 |
| EnteringCyt | *Salmonella* enters the cytosol | 0.004 |
| ProSalVac | Proliferation of *Salmonella* in the SCV | 0.000000411 |
| ProSalCyt | Proliferation of *Salmonella* in the cytosol | 0.000000595 |
| BeingCytIncapable | Cell becomes cytosolic-incapable | 650 |
| BeingCytCapable | Cell becomes cytosolic-capable | 350 |
| AdaptingSigCyt | Induces the adaptation of *Salmonella* to the cytosol | 1000 |
| AdaptingCyt | Increases the adaptability of *Salmonella* to the cytosol | 0.167 |
| DividingSalCyt | Increases the divisibility of *Salmonella* in the cytosol | 1000 |
| AdaptingSigVac | Induces the adaptation of *Salmonella* to the SCV | 1000 |
| AdaptingVac | Increases the adaptability of *Salmonella* to the SCV | 0.036 |
| DividingSalVac | Increases the divisibility of *Salmonella* in the SCV | 1000 |
| XenoDeg | Removes one bacterium from the cytosol via xenophagy | 0.00037 |
| XenoSigInducing | Induces the initiation and termination of xenophagy | 1000 |
| XenoIniCount | Increases the time counter of the initiation of xenophagy | 1000 |
| XenoDegInitiation | Activation of xenophagy | 0.56 |
| XenoEndCount | Increases the time counter of the termination of xenophagy | 1000 |
| XenoDegEnding | Termination of xenophagy | 0.069 |
| DeathCellCyt1 | Cell death due to a high number of cytosolic *Salmonella* | 1000 |
| DeathCellCyt2 | Removes cytosolic *Salmonella* in cytosolic-bacteria-induced cell death | 1000 |
| DeathCellVac1 | Cell death due to a high number of vacuolar *Salmonella* | 1000 |
| DeathCellVac2 | Removes vacuolar *Salmonella* in vacuolar-bacteria-induced cell death | 1000 |
| Death1 | Removes vacuolar *Salmonella* in cytosolic-bacteria-induced cell death | 1000 |
| Death2 | Removes cytosolic *Salmonella* in vacuolar-bacteria-induced cell death | 1000 |
| WashCount | Increases the time counter of the washing process | See Table 2 |
| WashInitiation | Generates a signal for the onset of the washing process | 1000 |
| Washing1 | Washing to remove non-invaded *Salmonella* in the infection medium | 1000 |
| Washing2 | Washing to remove non-invaded *Salmonella* in the infection medium | 1000 |
| Washing3 | Washing to remove non-invaded *Salmonella* on the cell surface | 1000 |

## References

[1] S. E. Majowicz, J. Musto, E. Scallan, F. J. Angulo, M. Kirk, S. J. O’Brien, T. F. Jones, A. Fazil, and R. M. Hoekstra, The Global Burden of Nontyphoidal *Salmonella* Gastroenteritis, Clin. Infect. Dis. 50, 882–889 (2010).

[2] L. A. Lee, N. D. Puhr, E. K. Maloney, N. H. Bean, and R. V. Tauxe, Increase in Antimicrobial-Resistant *Salmonella* Infections in the United States, 1989–1990, J. Infect. Dis. 170, 128–134 (1994).

[3] S. Kariuki, M. A. Gordon, N. Feasey, and C. M. Parry, Antimicrobial resistance and management of invasive *Salmonella* disease, Vaccine 33, C21–C29 (2015).

[4] M. E. Ohl and S. I. Miller, *Salmonella*: A Model for Bacterial Pathogenesis, Annu. Rev. Med. 52, 259–274 (2001).

[5] P. Garai, D. P. Gnanadhas, and D. Chakravortty, *Salmonella enterica* serovars Typhimurium and Typhi as model organisms: Revealing paradigm of host-pathogen interactions, Virulence 3, 377–388 (2012).

[6] A. Takeuchi, Electron microscope studies of experimental *Salmonella* infection. I. Penetration into the intestinal epithelium by *Salmonella* Typhimurium, Am. J. Pathol. 50, 109–136 (1967).

[7] A. Takeuchi and H. Sprinz, Electron-microscope studies of experimental *Salmonella* infection in the preconditioned guinea pig. II. Response of the intestinal mucosa to the invasion by *Salmonella* Typhimurium, Am. J. Pathol. 51, 137–161 (1967).

[8] J. E. Galán and R. Curtiss III, Cloning and molecular characterization of genes whose products allow *Salmonella typhimurium* to penetrate tissue culture cells, Proc. Natl. Acad. Sci. U. S. A. 86, 6383–6387 (1989).

[9] B. B. Finlay, S. Ruschkowski, and S. Dedhar, Cytoskeletal rearrangements accompanying *Salmonella* entry into epithelial cells, J. Cell Sci. 99, 283–296 (1991).

[10] F. Garcia-del Portillo, M. B. Zwick, K. Y. Leung, and B. B. Finlay, *Salmonella* induces the formation of filamentous structures containing lysosomal membrane glycoproteins in epithelial cells, Proc. Natl. Acad. Sci. U. S. A. 90, 10544–10548 (1993).

[11] F. Garcia-del Portillo and B. B. Finlay, Targeting of *Salmonella* typhimurium to Vesicles Containing Lysosomal Membrane Glycoproteins Bypasses Compartments with Mannose 6-Phosphate Receptors, J. Cell Biol. 129, 81–97 (1995).

[12] M. Hensel, J. E. Shea, C. Gleeson, M. D. Jones, E. Dalton, and D. W. Holden, Simultaneous Identification of Bacterial Virulence Genes by Negative Selection, Science 269, 400–403 (1995).

[13] J. E. Shea, M. Hensel, C. Gleeson, and D. W. Holden, Identification of a virulence locus encoding a second type III secretion system in *Salmonella* typhimurium, Proc. Natl. Acad. Sci. U. S. A. 93, 2593– 2597 (1996).

[14] J. E. Galán, Molecular genetic bases of *Salmonella* entry into host cells, Mol. Microbiol. 20, 263–271 (1996).

[15] E. A. Groisman and H. Ochman, How *Salmonella* became a pathogen, Trends Microbiol. 5, 343–349 (1997).

[16] C. M. Collazo and J. E. Galán, The invasion-associated type III system of *Salmonella* Typhimurium directs the translocation of Sip proteins into the host cell, Mol. Microbiol. 24, 747–756 (1997).

[17] J. H. Brumell, O. Steele-Mortimer, and B. B. Finlay, Bacterial invasion: Force feeding by *Salmonella*, Curr. Biol. 9, R277–R280 (1999).

[18] K. Eichelberg and J. E. Galán, Differential Regulation of *Salmonella* Typhimurium Type III Secreted Proteins by Pathogenicity Island 1 (SPI-1)-Encoded Transcriptional Activators InvF and HilA, Infect. Immun. 67, 4099–4105 (1999).

[19] B. B. Finlay and J. H. Brumell, *Salmonella* interactions with host cells: *in vitro* to *in vivo*, Philos. Trans. R. Soc. Lond. B Biol. Sci. 355, 623–631 (2000).

[20] C. R. Beuzón, S. Méresse, K. E. Unsworth, J. Ruíz-Albert, S. Garvis, S. R. Waterman, T. A. Ryder, E. Boucrot, and D. W. Holden, *Salmonella* maintains the integrity of its intracellular vacuole through the action of SifA, EMBO J. 19, 3235–3249 (2000).

[21] J. E. Galán and D. Zhou, Striking a balance: Modulation of the actin cytoskeleton by *Salmonella*, Proc. Natl. Acad. Sci. U. S. A. 97, 8754– 8761 (2000).

[22] D. Zhou, L.-M. Chen, L. Hernandez, S. B. Shears, and J. E. Galán, A *Salmonella* inositol polyphosphatase acts in conjunction with other bacterial effectors to promote host cell actin cytoskeleton rearrangements and bacterial internalization, Mol. Microbiol. 39, 248–259 (2001).

[23] I. Hansen-Wester and M. Hensel, *Salmonella* pathogenicity islands encoding type III secretion systems, Microbes Infect. 3, 549–559 (2001).

[24] D. Zhou and J. Galán, *Salmonella* entry into host cells: the work in concert of type III secreted effector proteins, Microbes Infect. 3, 1293– 1298 (2001).

[25] J. E. Galán, *Salmonella* Interactions with Host Cells: Type III Secretion at Work, Annu. Rev. Cell Dev. Biol. 17, 53–86 (2001).

[26] O. Steele-Mortimer, J. H. Brumell, L. A. Knodler, S. Méresse, A. Lopez, and B. B. Finlay, The invasion-associated type III secretion system of *Salmonella enterica* serovar Typhimurium is necessary for intracellular proliferation and vacuole biogenesis in epithelial cells, Cell. Microbiol. 4, 43–54 (2002).

[27] J. H. Brumell, P. Tang, M. L. Zaharik, and B. B. Finlay, Disruption of the *Salmonella*-Containing Vacuole Leads to Increased Replication of *Salmonella enterica* Serovar Typhimurium in the Cytosol of Epithelial Cells, Infect. Immun. 70, 3264–3270 (2002).

[28] C. R. Beuzón, S. P. Salcedo, and D. W. Holden, Growth and killing of a *Salmonella enterica* serovar Typhimurium *sifA* mutant strain in the cytosol of different host cell lines, Microbiol. 148, 2705–2715 (2002).

[29] L. A. Knodler and O. Steele-Mortimer, Taking Possession: Biogenesis of the *Salmonella*-Containing Vacuole, Traffic 4, 587–599 (2003).

[30] A. J. Perrin, X. Jiang, C. L. Birmingham, N. S. Y. So, and J. H. Brumell, Recognition of Bacteria in the Cytosol of Mammalian Cells by the Ubiquitin System, Curr. Biol. 14, 806–811 (2004).

[31] C. Altier, Genetic and Environmental Control of *Salmonella* Invasion, J. Microbiol. 43, 85–92 (2005).

[32] C. L. Birmingham, A. C. Smith, M. A. Bakowski, T. Yoshimori, and J. H. Brumell, Autophagy Controls *Salmonella* Infection in Response to Damage to the *Salmonella*-containing Vacuole, J. Biol. Chem. 281, 11374–11383 (2006).

[33] B. Coburn, G. A. Grassl, and B. B. Finlay, *Salmonella*, the host and disease: a brief review, Immunol. Cell Biol. 85, 112–118 (2007).

[34] D. Drecktrah, L. A. Knodler, D. Howe, and O. Steele-Mortimer, *Salmonella* Trafficking is Defined by Continuous Dynamic Interactions with the Endolysosomal System, Traffic 8, 212–225 (2007).

[35] A. Orvedahl and B. Levine, Eating the enemy within: autophagy in infectious diseases, Cell Death Differ. 16, 57–69 (2009).

[36] E. J. McGhie, L. C. Brawn, P. J. Hume, D. Humphreys, and V. Koronakis, *Salmonella* takes control: effector-driven manipulation of the host, Curr. Opin. Microbiol. 12, 117–124 (2009).

[37] K. Ray, B. Marteyn, P. J. Sansonetti, and C. M. Tang, Life on the inside: the intracellular lifestyle of cytosolic bacteria, Nat. Rev. Microbiol. 7, 333–340 (2009).

[38] T. L. M. Thurston, G. Ryzhakov, S. Bloor, N. von Muhlinen, and F. Randow, The TBK1 adaptor and autophagy receptor NDP52 restricts the proliferation of ubiquitin-coated bacteria, Nat. Immun. 10, 1215–1221 (2009).

[39] L. A. Knodler, B. A. Vallance, J. Celli, S. Winfree, B. Hansen, M. Montero, and O. Steele-Mortimer, Dissemination of invasive *Salmonella* via bacterial-induced extrusion of mucosal epithelia, Proc. Natl. Acad. Sci. U. S. A. 107, 17733–17738 (2010).

[40] B. Misselwitz, S. K. Kreibich, S. Rout, B. Stecher, B. Periaswamy, and W.-D. Hardt, *Salmonella enterica* Serovar Typhimurium Binds to HeLa Cells via Fim-Mediated Reversible Adhesion and Irreversible Type Three Secretion System 1-Mediated Docking, Infect. Immun. 79, 330–341 (2011).

[41] B. Misselwitz, S. Dilling, P. Vonaesch, R. Sacher, B. Snijder, M. Schlumberger, S. Rout, M. Stark, C. von Mering, L. Pelkmans, and W.-D. Hardt, RNAi screen of *Salmonella* invasion shows role of COPI in membrane targeting of cholesterol and Cdc42, Mol. Syst. Biol. 7, 474 (2011).

[42] P. Wild, H. Farhan, D. G. McEwan, S. Wagner, V. V. Rogov, N. R. Brady, B. Richter, J. Korac, O. Waidmann, C. Choudhary, V. Dötsch, D. Bumann, and I. Dikic, Phosphorylation of the Autophagy Receptor Optineurin Restricts *Salmonella* Growth, Science 333, 228–233 (2011).

[43] L. A. Knodler and J. Celli, Eating the strangers within: host control of intracellular bacteria via xenophagy, Cell. Microbiol. 13, 1319–1327 (2011).

[44] A. J. Müller, P. Kaiser, K. E. J. Dittmar, T. C. Weber, S. Haueter, K. Endt, P. Songhet, C. Zellweger, M. Kremer, H.-J. Fehling, and W.-D. Hardt, *Salmonella* Gut Invasion Involves TTSS-2-Dependent Epithelial Traversal, Basolateral Exit, and Uptake by Epithelium-Sampling Lamina Propria Phagocytes, Cell Host Microbe 11, 19–32 (2012).

[45] T. Dandekar, A. Fieselmann, J. Popp, and M. Hensel, *Salmonella enterica*: a surprisingly well-adapted intracellular lifestyle, Front. Microbiol. 3, 164 (2012).

[46] P. Malik-Kale, S. Winfree, and O. Steele-Mortimer, The Bimodal Lifestyle of Intracellular *Salmonella* in Epithelial Cells: Replication in the Cytosol Obscures Defects in Vacuolar Replication, PLoS ONE 7, e38732 (2012).

[47] I. Tattoli, M. T. Sorbara, D. Vuckovic, A. Ling, F. Soares, L. A. M. Carneiro, C. Yang, A. Emili, D. J. Philpott, and S. E. Girardin, Amino Acid Starvation Induced by Invasive Bacterial Pathogens Triggers an Innate Host Defense Program, Cell Host Microbe 11, 563–575 (2012).

[48] B. Misselwitz, N. Barrett, S. Kreibich, P. Vonaesch, D. Andritschke, S. Rout, K. Weidner, M. Sormaz, P. Songhet, P. Horvath, M. Chabria, V. Vogel, D. M. Spori, P. Jenny, and W.-D. Hardt, Near Surface Swimming of *Salmonella* Typhimurium Explains Target-Site Selection and Cooperative Invasion, PLoS Pathog. 8, e1002810 (2012).

[49] P. Velge, A. Wiedemann, M. Rosselin, N. Abed, Z. Boumart, M. Chaussé, O. Grépinet, F. Namdari, S. M. Roche, A. Rossignol, and I. Virlogeux-Payant, Multiplicity of *Salmonella* entry mechanisms, a new paradigm for *Salmonella* pathogenesis, MicrobiologyOpen 1, 243–258 (2012).

[50] I. Tattoli, D. J. Philpott, and S. E. Girardin, The bacterial and cellular determinants controlling the recruitment of mTOR to the *Salmonella*- containing vacuole, Biol. Open 1, 1215–1225 (2012).

[51] T. P. Moest and S. Méresse, *Salmonella* T3SSs: successful mission of the secret(ion) agents, Curr. Opin. Microbiol. 16, 38–44 (2013).

[52] A. Fàbrega and J. Vila, *Salmonella enterica* Serovar Typhimurium Skills To Succeed in the Host: Virulence and Regulation, Clin. Microbiol. Rev. 26, 308–341 (2013).

[53] E.-k. Jo, J.-M. Yuk, D.-M. Shin, and C. Sasakawa, Roles of autophagy in elimination of intracellular bacterial pathogens, Front. Immunol. 4, 97 (2013).

[54] J. L. Benjamin, R. Sumpter Jr., B. Levine, and L. V. Hooper, Intestinal Epithelial Autophagy Is Essential for Host Defense against Invasive Bacteria, Cell Host Microbe 13, 723–734 (2013).

[55] L. A. Knodler, V. Nair, and O. Steele-Mortimer, Quantitative Assessment of Cytosolic *Salmonella* in Epithelial Cells, PLoS ONE 9, e84681 (2014).

[56] J. Huang and J. H. Brumell, Bacteria-autophagy interplay: a battle for survival, Nat. Rev. Microbiol. 12, 101–114 (2014).

[57] M. Lorkowski, A. Felipe-López, C. A. Danzer, N. Hansmeier, and M. Hensel, *Salmonella enterica* Invasion of Polarized Epithelial Cells Is a Highly Cooperative Effort, Infect. Immun. 82, 2657–2667 (2014).

[58] L. A. Knodler, *Salmonella enterica*: living a double life in epithelial cells, Curr. Opin. Microbiol. 23, 23–31 (2015).

[59] M. Wrande, H. Andrews-Polymenis, D. J. Twedt, O. Steele-Mortimer, S. Porwollik, M. McClelland, and L. A. Knodler, Genetic Determinants of *Salmonella enterica* Serovar Typhimurium Proliferation in the Cytosol of Epithelial Cells, Infect. Immun. 84, 3517–3526 (2016).

[60] S. Castanheira and F. García-del Portillo, *Salmonella* Populations inside Host Cells, Front. Cell. Infect. Microbiol. 7, 432 (2017).

[61] R. Johnson, E. Mylona, and G. Frankel, Typhoidal *Salmonella*: Distinctive virulence factors and pathogenesis, Cell. Microbiol. 20, e12939 (2018).

[62] S. Wu, Y. Shen, S. Zhang, Y. Xiao, and S. Shi, *Salmonella* Interacts With Autophagy to Offense or Defense, Front. Microbiol. 11, 721 (2020).

[63] S. A. Fattinger, D. Böck, M. L. Di Martino, S. Deuring, P. Samperio Ventayol, V. Ek, M. Furter, S. Kreibich, F. Bosia, A. A. Müller-xsHauser, B. D. Nguyen, M. Rohde, M. Pilhofer, W.-D. Hardt, and M. E. Sellin, *Salmonella* Typhimurium discreet-invasion of the murine gut absorptive epithelium, PLoS Pathog. 16, e1008503 (2020).

[64] S. J. Taylor and S. E. Winter, *Salmonella* finds a way: Metabolic versatility of *Salmonella enterica* serovar Typhimurium in diverse host environments, PLoS Pathog. 16, e1008540 (2020).

[65] A. Kehl, J. Noster, and M. Hensel, Eat in or Take out? Metabolism of Intracellular *Salmonella enterica*, Trends Microbiol. 28, 644–654 (2020).

[66] H. Bao, S. Wang, J.-H. Zhao, and S.-L. Liu, *Salmonella* secretion systems: Differential roles in pathogen-host interactions, Microbiol. Res. 241, 126591 (2020).

[67] P. Geiser, M. L. Di Martino, P. Samperio Ventayol, J. Eriksson, E. Sima, A. K. Al-Saffar, D. Ahl, M. Phillipson, D.-L. Webb, M. Sundbom, P. M. Hellström, and M. E. Sellin, *Salmonella enterica* Serovar Typhimurium Exploits Cycling through Epithelial Cells To Colonize Human and Murine Enteroids, mBio 12, e02684–20 (2021).

[68] J. E. Galán, *Salmonella* Typhimurium and inflammation: a pathogen-centric affair, Nat. Rev. Microbiol. 19, 716–725 (2021).

69. [68] B. A. Chong, K. G. Cooper, L. Kari, O. R. Nilsson, C. Hillman, A. Fleming, Q. Wang, V. Nair, and O. Steele-Mortimer, Cytosolic replication in epithelial cells fuels intestinal expansion and chronic fecal shedding of *Salmonella* Typhimurium, Cell Host Microbe 29, 1177–1185 (2021).

[70] T. R. Powers, A. L. Haeberle, A. V. Predeus, D. L. Hammarlöf, J. A. Cundiff, Z. Saldaña-Ahuactzi, K. Hokamp, J. C. D. Hinton, and L. A. Knodler, Intracellular niche-specific profiling reveals transcriptional adaptations required for the cytosolic lifestyle of *Salmonella enterica*, PLoS Pathog. 17, e1009280 (2021).

[71] J. Hannig, Application and method development in computational systems biology: Petri nets to study knockouts and dynamics of Salmonella infection, Ph. D. dissertation (Goethe University Frankfurt, Frankfurt am Main, Germany, 2018).

[72] C. Das, A. Dutta, H. Rajasingh, and S. S. Mande, Understanding the sequential activation of Type III and Type VI Secretion Systems in *Salmonella* typhimurium using Boolean modeling, Gut Pathog. 5, 28 (2013).

[73] J. Scheidel, L. Amstein, J. Ackermann, I. Dikic, and I. Koch, *In Silico* Knockout Studies of Xenophagic Capturing of *Salmonella*, PLoS Comput. Biol. 12, e1005200 (2016).

[74] Y. Chen, G. Wang, H. Cai, Y. Sun, L. Quyang, and B. Liu, Deciphering the Rules of in Silico Autophagy Methods for Expediting Medicinal Research, J. Med. Chem. 62, 6831–6842 (2019).

[75] E. Fiskin, T. Bionda, I. Dikic, and C. Behrends, Global Analysis of Host and Bacterial Ubiquitinome in Response to *Salmonella* Typhimurium Infection, Mol. Cell 62, 967–981 (2016).

[76] R. W. Kasinskas and N. S. Forbes, *Salmonella* typhimurium Specifically Chemotax and Proliferate in Heterogeneous Tumor Tissue in Vitro, Biotechnol. Bioeng. 94, 710–721 (2006).

[77] S. P. Brown, S. J. Cornell, M. Sheppard, A. J. Grant, D. J. Maskell, B. T. Grenfell, and P. Mastroeni, Intracellular Demography and the Dynamics of *Salmonella enterica* Infections, PLoS Biol. 4, e349 (2006).

[78] S. C. Andrés, L. Giannuzzi, and N. E. Zaritzky, Mathematical Modeling of Microbial Growth in Packaged Refrigerated Orange Juice Treated with Chemical Preservatives, J. Food Sci. 66, 724–728 (2001).

[79] V. K. Juneja, M. V. Melendres, L. Huang, J. Subbiah, and H. Thippareddi, Mathematical modeling of growth of *Salmonella* in raw ground beef under isothermal conditions from 10 to 45 °C, Int. J. Food Microbiol. 131, 106–111 (2009).

[80] W. Pan and D. W. Schaffner, Modeling the Growth of *Salmonella* in Cut Red Round Tomatoes as a Function of Temperature, J. Food Prot. 73, 1502–1505 (2010).

[81] P. R. Velugoti, L. K. Bohra, V. K. Juneja, L. Huang, A. L. Wesseling, J. Subbiah, and H. Thippareddi, Dynamic model for predicting growth of *Salmonella* spp. in ground sterile pork, Food Microbiol. 28, 796–803 (2011).

[82] K. McDonald and D.-W. Sun, Predictive food microbiology for the meat industry: a review, Int. J. Food Microbiol. 52, 1–27 (1999).

[83] T. A. McMeekin, J. Olley, D. A. Ratkowsky, and T. Ross, Predictive microbiology: towards the interface and beyond, Int. J. Food Microbiol. 73, 395–407 (2002).

[84] P. P. Chapagain, J. S. van Kessel, J. S. Karns, D. R. Wolfgang, E. Hovingh, K. A. Nelen, Y. H. Schukken, and Y. T. Grohn, A mathematical model of the dynamics of *Salmonella* Cerro infection in a US dairy herd, Epidemiol. Infect. 136, 263–272 (2008).

[85] Y. Xiao, D. Clancy, N. P. French, and R. G. Bowers, A semi-stochastic model for *Salmonella* infection in a multi-group herd, Math. Biosci. 200, 214–233 (2006).

[86] A. D. C. Berriman, D. Clancy, H. E. Clough, and R. M. Christley, Semi-stochastic models for *Salmonella* infection within finishing pig units in the UK, Math. Biosci. 245, 148–156 (2013).

[87] A. J. Grant, O. Restif, T. J. McKinley, M. Sheppard, D. J. Maskell, and P. Mastroeni, Modelling within-Host Spatiotemporal Dynamics of Invasive Bacterial Disease, PLoS Biol. 6, e74 (2008).

[88] J. R. Gog, A. Murcia, N. Osterman, O. Restif, T. J. McKinley, M. Sheppard, S. Achouri, B. Wei, P. Mastroeni, J. L. N. Wood, D. J. Maskell, P. Cicuta, and C. E. Bryant, Dynamics of *Salmonella* infection of macrophages at the single cell level, J. R. Soc. Interface 9, 2696–2707 (2012).

[89] D. T. Gillespie, A General Method for Numerically Simulating the Stochastic Time Evolution of Coupled Chemical Reactions, J. Comput. Phys. 22, 403–434 (1976).

[90] D. T. Gillespie, Exact Stochastic Simulation of Coupled Chemical Reactions, J. Phys. Chem. 81, 2340–2361 (1977).

[91] T. Székely Jr. and K. Burrage, Stochastic simulation in systems biology, Comput. Struct. Biotechnol. J. 12, 14–25 (2014).

[92] D. A. Kennedy, V. Dukic, and G. Dwyer, Pathogen Growth in Insect Hosts: Inferring the Importance of Different Mechanisms Using Stochastic Models and Response-Time Data, Am. Nat. 184, 407–423 (2014).

[93] C. Chaouiya, Petri net modelling of biological networks, Brief. Bioinform. 8, 210–219 (2007).

[94] I. Koch, W. Reisig, and F. Schreiber, Modeling in Systems Biology: The Petri Net Approach (Springer, London, 2011).

[95] T. Dunn, Image Segmentation and Modelling of Host-Pathogen Dynamics of Salmonella, M. Sc. thesis (Dalhousie University, Nova Scotia, Canada, 2016).

[96] M. Madigan, J. Martinko, K. Bender, D. Buckley, and D. Stahl, Brock Biology of Microorganisms, 14th ed. (Pearson Education Limited, Harlow, United Kingdom, 2015).

[97] P. Balazki, K. Lindauer, J. Einloft, J. Ackermann, and I. Koch, MONALISA for stochastic simulations of Petri net models of biochemical systems, BMC Bioinformatics 16, 215 (2015).

[98] L. Zhao, C. D. Kroenke, J. Song, D. Piwnica-Worms, J. J. H. Ackerman, and J. J. Neil, Intracellular water-specific MR of microbead-adherent cells: the HeLa cell intracellular water exchange lifetime, NMR Biomed. 21, 159–164 (2008).

[99] C. Pramoolsinsap and V. Viranuvatti, *Salmonella* hepatitis, J. Gastroenterol. Hepatol. 13, 745–750 (1998).

[100] A. Vazquez-Torres, J. Jones-Carson, A. J. Bäumler, S. Falkow, R. Valdivia, W. Brown, M. Le, R. Berggren, W. T. Parks, and F. C. Fang, Extraintestinal dissemination of *Salmonella* by CD18-expressing phagocytes, Nature 401, 804–808 (1999).

[101] K. L. Rosche, A. T. Aljasham, J. N. Kipfer, B. T. Piatkowski, and V. Konjufca, Infection with *Salmonella enterica* Serovar Typhimurium Leads to Increased Proportions of F4/80^+^ Red Pulp Macrophages and Decreased Proportions of B and T Lymphocytes in the Spleen, PLoS ONE 10, e0130092 (2015).

[102] G. Gonzalez-Escobedo, J. M. Marshall, and J. S. Gunn, Chronic and acute infection of the gall bladder by *Salmonella* Typhi: understanding the carrier state, Nat. Rev. Microbiol. 9, 9–14 (2011).

[103] N. M. van Sorge, P. A. Zialcita, S. H. Browne, D. Quach, D. G. Guiney, and K. S. Doran, Penetration and Activation of Brain Endothelium by *Salmonella enterica* Serovar Typhimurium, J. Infect. Dis. 203, 401–405 (2011).

[104] H. Xin, Y. Li, D. Xu, Y. Zhang, C.-H. Chen, and B. Li, Single Upconversion Nanoparticle-Bacterium Cotrapping for Single-Bacterium Labeling and Analysis, Small 13, 1603418 (2017).

